# Incorporation of complex narratives into dreaming

**DOI:** 10.1101/2025.02.11.637675

**Authors:** Jessica Palmieri, Deniz Kumral, Sammy-Jo Wymer, Susanne Kirchner, Maximilian Schneider, Steffen Gais, Monika Schönauer

## Abstract

Reactivation of waking neuronal activity during sleep holds a functional role in memory consolidation. Reprocessing of daytime memory in dreams might aid later memory performance in a similar way. Numerous findings hint at a link between dreaming and sleep-dependent memory processing, however, studies investigating day-residue incorporation in dreaming led to mixed results so far. In this study, we used a naturalistic learning paradigm aimed at biasing dream content by manipulating pre-sleep experience. Participants listened to one of four different audiobooks while falling asleep and were awoken several times during the night to report their dreams. Afterwards, we tested how well they remembered the content of the audiobook. We then asked three blind raters to guess, based solely on anonymized dream reports, which audiobook someone had listened to before experiencing a dream. Our findings show that dreams across the whole night and from both NREM and REM awakenings contain specific information about the content of narratives studied before sleep. Moreover, participants with stronger incorporation of the audiobook in their dreams showed a tendency to recognize more audiobook content across the sleep period. Together, these findings suggest that salient day-time experiences resurface in dreams and that content selected for consolidation during sleep is more strongly incorporated.

## INTRODUCTION

We are asleep for approximately one third of our lives. During this time, we are cut off from the external world, yet experience vivid internal mentation, known as dreams. Generations have pondered over the meaning of this conscious experience during sleep [1,2]. It has long been known that dreaming is somehow linked to daytime experience. Freud, who studied dreams intensively, hence termed the phrase “daytime residue” [3]. Several authors refer to the correspondence between waking life and dream content as the “continuity hypothesis” [4–8], according to which salient elements, thoughts, or concerns of one’s waking experience are re-expressed in dreams. Empirical findings supported this view, as it has been shown that relevant aspects of waking life [9], as well as significant life events [10,11], affect dream content. Recent research has thus started to address *why* and *how* our real-world experience becomes oneiric material.

### Dreaming as an indication of memory reprocessing

A number of findings suggests that dream content might be linked to sleep-dependent memory consolidation, possibly reflecting a conscious manifestation of the neural processes that underlie nighttime memory processing [12–14]. For one, neuronal activation during sleep is directly linked to previous waking experience in animals, such that cells which were active during exploration of an environment also show stronger activity during later sleep [15,16], and fire in the same sequences as while the animal was moving about [17]. More recently, spontaneous reactivation of learning-related brain activity patterns has been reported in humans and was successfully related to subsequent memory for the studied material [18–22].

Accordingly, dreaming at least partially reflects novel learning. When participants played the game Tetris for extended periods before falling asleep, Tetris-related imagery was observed in reports of hypnagogic images in participants woken from the sleep onset period [23]. Kusse and colleagues [24] independently replicated this impact of the game Tetris on sleep onset imagery. In another study employing a downhill skiing arcade game before sleep, 30% of sleep onset reports showed direct incorporation of the game [25]. Similar findings were observed in studies examining the relationship between the incorporation of pre-sleep experiences into dreams during more consolidated sleep, and later memory performance [26–28]. Participants who dreamt of a spatial navigation task were better at performing the task after sleep [26,29]. Moreover, if the content of participants’ dreams was related to the category of previously studied word-picture associations, they retained these memories better across the night [27]. In an episodic memory task using odors and landscapes, Plailyl et al. [28] showed that dream incorporation of task-related content improved visuo-spatial memory for the implicitly encoded episodes. Note, however, that other studies investigating the relationship between dream incorporation and later memory performance yielded mixed results [28,30–36] (see [28] for a comprehensive summary).

Shared properties of neural and mental reactivation corroborate the idea of a connection between the two: At the neural level, reactivation events are not precisely identical to the actual real-world encounter [37], they occur at a different time scale [38], in a fragmented fashion, through discrete bursts of neuronal activity [39]. This suggests that memory reinstatement arises through the activation of discrete episodic units of information. In dreaming, conscious experiences with relatable pre-sleep content do not replay waking events exactly as they occur. When an incorporation of wake-life events is observed, it is rarely the faithful reproduction of an episodic memory in its entireness [40–42].

It is unclear whether content incorporation in dreaming is regulated by the same neural mechanisms across all sleep phases. For instance, sleep onset incorporations seem to occur independently of the hippocampal system: Patients with extensive medial temporal lobe damage including the hippocampus, with no memory for the pre-sleep game session, could still report Tetris blocks-related imagery during sleep onset [23]. On the contrary, an intact hippocampus seems to be necessary for typical consolidated sensory-rich dreaming to occur, as well as for day-residue elaborations in dreams to appear [43–45]. This suggests that sleep onset mentation and more consolidated dreaming, and consequently the incorporation of pre-sleep elements during both phenomena, might partly rely on distinct neural mechanisms. More generally, the access to previous memory as a source of dream construction might drastically vary depending on brain connectivity and sleep state. Effective communication across memory systems strongly depends on a finely regulated milieu of neurotransmitters that shifts across sleep stages [46].

### NREM and REM dreaming

While dreaming was initially attributed exclusively to REM sleep [47,48], dream reports can be collected from all sleep stages and across the whole night [1,12,49–54]. Dreaming has been observed in up to 50% of cases from NREM sleep awakenings, especially from sleep stage 2 [49,55,56]. Moreover, evidence showed that pharmacological REM sleep suppression does not significantly affect dream recall [57,58]. Various studies, however, reported both quantitative and qualitative differences between dream content deriving from NREM and REM awakenings. Dreaming from NREM sleep seems to be more thought-like and conceptual [59,60], whereas REM dreaming is usually described as more story-like, hallucinatory, and bizarre [61,62], and is usually associated with longer reports [63,64,53]. The distinct neurophysiological features characterizing NREM and REM sleep might explain the observed differences between REM and NREM dreaming, which also manifests in the way memory consolidation differently unfolds in the two sleep stages [65]. Studies showed that early-night dreams-when NREM is predominant and ACh levels are at their lowest-more frequently incorporate recent experiences compared to late-night dreams [66,67], and tend to be more episodic rather than semantic in nature [68]. Dream construction later in the night, especially during REM sleep, might instead mirror cortical networks re-organization, with dream contents thus being more associative in nature [68–70], and displaying more frequent semantic references [12,53]. However, the existing literature is currently not unanimous on the distinct features of NREM and REM dreaming [56,58,60], as well as their respective memory sources or incorporation dynamics across the night [53,60,71]. After combining and re-analyzing previous data, Baylor & Cavallero [40], in contrast with Cavallero et al.[53], observed higher episodic sources in dream reports from NREM compared to REM awakenings, but could not find any stage effect for semantic memory sources. In a recent study, a higher number of mixed memory sources have been observed in both N1 and REM dreams compared to consolidated NREM dreams [67]. Importantly, ultradian cycles and other chronobiological factors also contribute to the quality of dreams and memory sources selection [72–74]. Recent evidence suggests that time of the night, more than sleep stages alone, determines which memory sources more likely will contribute to dream construction across sleep periods [67]. For a comprehensive understanding of the accessibility of memory sources in dreams throughout the night, the interplay between all the above-mentioned may thus be relevant.

### Incorporation of the experimental situation into dreams

In studies on dreaming, it is common to observe dream incorporation of elements related to the experimental situation [75]. Examples of recurrent themes can be the presence of the experimenter, sleeping in the laboratory, having EEG electrodes or cables attached, and being examined. Explicit references to the experiment were found in 6-32% of dreams across different studies, with the proportion rising to 32% - 68% when considering more implicit laboratory references as well [6,76] (for more studies see [75]). When present, this phenomenon may occur at the expense of the experimental manipulation itself: In early studies, pre-sleep stimuli incorporation was detected in only 5-8% of reports [77,78]. Because many argue that salient and engaging content is what mostly gets incorporated into dreams [14], the experience of sleeping in a laboratory environment while wearing an EEG device might turn out to be more notable and novel to participants than the to-be studied content. In a recent study, Picard-Deland and Nielsen [79] observed lab-related incorporation in over a third of all dreams, in almost half of participants who recalled a dream. Most experimental setting incorporations in Picard-Deland & Nielsen study were rated as prospective. This would further support dreaming’s relation with reactivation mechanisms and memory processes: Neuronal activation seems to mostly prioritize goal-relevant, salient memories [80], such that consolidation would be particularly adaptive for immediate-or distant-future use [81].

### Aims of this study

In the current study, we investigated whether conscious experience during sleep reflects memory reprocessing using naturalistic stimulus material. We hypothesized that memory content encoded before going to bed should resurface in the conscious experience of participants during subsequent sleep. If it is possible to determine the content of a previous learning episode based on dream reports alone, they must contain information highly specific to this experience. To test this, we had participants listen to one of four different audiobooks while falling asleep. They were asked to remember the content well enough so they could retrieve it later. Participants were awoken several times during the night and asked to report their dreams. Afterwards, we tested how well they remembered the content of the audiobook. This procedure was repeated multiple times throughout the night. We then asked three blind raters to guess, based solely on anonymized dream reports, which audiobook someone had listened to before experiencing a dream. We further had them indicate how much information about the audiobook they detected in the dream report. Our results show that dreams across the whole night and from both NREM and REM awakenings contain specific information about the content of narratives studied before sleep. Moreover, participants with stronger incorporation of the audiobook in their dreams showed a tendency to recognize the audiobook content better across the sleep period. Finally, we also examined whether the incorporation of the experimental situation into dreams interferes with incorporation of the learning material.

## METHODS

### Participants

20 subjects (10 male) aged 20 – 30 (25.5 ± 2.7 [mean ± s.d.]) participated in this study. One participant did not report any dream throughout the night and was thus excluded from all analyses (remaining N = 19).

Participants were healthy, non-smokers, and did not ingest any alcohol, caffeine or medication other than oral contraceptives on the days of the experiment. The participants reported sleeping between 6 and 10 hours per night, had a regular circadian rhythm, and were not extreme morning or evening chronotypes, measured by the Munich Chronotype Questionnaire (MCTQ) [82]. They had no shift work or long-distance flights during the six weeks preceding the experiment and did not have any sleep-related pathology. All participants were right-handed, as confirmed by the Edinburgh Handedness questionnaire [83]. Only participants who reported to frequently recall dreams (at least three dreams per week) were selected for this study. The experiment was approved by the local ethics committee (Department of Psychology, Ludwig-Maximilians-Universität München). Informed consent was obtained from all subjects.

### Experimental Procedure

All participants visited our laboratory twice, once for an adaptation night to become familiar with the experimental procedure and environment, and again for the night of the actual experiment. Subjects had to indicate beforehand whether they had read or watched the movies of the following books: Cornelia Funke – Inkheart, Agatha Christie – The Mystery of the Blue Train, Daniel Kehlmann – Measuring the World, Rick Riordan – Percy Jackson & the Olympians: The Lightning Thief. None of the participants admitted to the study were familiar with the audiobook content. On the day of the experiment, participants fell asleep while listening to one of the four audiobooks. The books were randomly assigned to the participants. They were instructed that they should listen closely, such that they would be able to report what occurred in the audiobook passage upon awakening, but to fall asleep when they felt the need. At the same time, a tone played every 15 seconds. Participants were asked to press any key on a computer mouse in response to the tone to confirm they were paying attention. The audiobook was turned off by the experimenter once they reached consolidated stage 2 sleep and stopped responding to the tone. The audio was played through external speakers at a volume of 45 dB ± 2.92 dB (mean ± SD), measured using a sound meter at the approximate location of the participants’ head in bed, ensuring the story was audible. From the moment participants were given the opportunity to fall asleep, they were awoken by the experimenter entering their room and addressing them by name after approximately one cycle (87 ± 16 min [mean ± SD]). If that was not sufficient to rouse the participant, the experimenter touched them lightly on the arm while addressing them again. Participants were informed of this procedure. Upon awakening, they were questioned about their conscious experience immediately before waking up (collection of dream reports, see below for details) and subsequently asked to retrieve the audiobook passage they had listened to before falling asleep (memory retrieval, see below for details). They were then instructed to go back to sleep, resuming to listen to the same audiobook at the point where they had left off before falling asleep, again paying close attention such that they could report what happened in the audiobook upon awakening. This procedure was repeated up to five times per night. The audiobook was played from beginning to end. Participants listened to an average of 35 ± 28 min [mean ± SD] minutes of audiobook before falling asleep. Two participants finished their audiobook before the experimental night was over. They continued the experiment listening to a different, additional audiobook, otherwise following the same procedure as reported above.

### Dream reports

After each sleep period, participants were questioned about their conscious experience at the moment of awakening. Conscious experience at awakening is considered to reflect the content of the dream that is experienced during sleep immediately before [84]. Dream reports were collected in a standardized manner by asking participants repeatedly, up to three times, what went through their mind immediately before waking up, and to report any conscious experience they might have had. If the participants were able to remember any conscious experience from the preceding sleep period, we instructed them that they should proceed to give a detailed report on who participated in the dream, where the dream was set, and what happened in the dream. We recorded their full dream report, asking up to three times “Can you recall more?” We then proceeded to inquire further details regarding the dream content with a standardized dream questionnaire: Who had been part of the dream? Where had the dream taken place? What happened in the dream? What was the perspective of the dream? Did the participant experience any emotions while dreaming? What would they name as the central element of their dream? Only questions that were predefined in this questionnaire were used to assess the content of the participant’s dream. All participants had to answer all questions, but had the option to indicate that they had no memory for any particular question. Questions were always asked in the same standardized order.

### Memory retrieval

After reporting their dreams, participants had to retrieve the audiobook content. They were first asked to freely recall what happened in the audiobook passage they had listened to before going to bed. To assess recognition memory, participants were then aurally presented with segments of the audiobook they had previously listened to and asked to indicate, for individual sentences, whether they remembered having listened to it earlier (*certainly yes, do not know, certainly not*). The whole questioning protocol was recorded on tape and later transcribed by someone not involved in any other part of the experimental procedure. The transcribed text of the free memory recall was then redacted to retain only content relevant to participants’ memory of the audiobook. Specifically, side remarks such as “what a strange name”, statements like “I do not know”, “I cannot remember”, “you know what I mean”, as well as repetitions and filler words (e.g “uhmm”) were removed. The remaining text was then further processed. Simple active language was used to eliminate individual language idiosyncrasies and to ensure that transcript length, relative to informational content, was as comparable as possible across participants. Free recall performance was assessed as number of words in the free recall divided by the time (in minutes) listened to the audiobook while participants were awake. Recognition performance was assessed as the audiobook runtime of yes responses divided by the audiobook runtime while awake. Note that no further measures were implemented to assess the accuracy of the recalled content. After the dream recall and audiobook retrieval, participants fell asleep while listening to the subsequent part of the same audiobook. All participants listened to at least 5 minutes of the audiobook before falling asleep (ranging from 6 to 121 minutes).

### Polysomnography

Sleep EEG was recorded using a 128-channel active Ag/AgCl electrode system (ActiCap, Brain Products, Gilching, Germany) with a sampling rate of 1 kHz and a high-pass filter at 0.1 Hz. Electrode placement followed the extended international 10–20 system [85]. For sleep staging, the recordings were segmented into 30-second epochs, and linear derivations were computed (EEG: C3 and C4 referenced to the contralateral mastoids, vertical and horizontal eye movements, and EMG activity). Sleep stages were assessed using electrodes C3 and C4 based on standard criteria, with two independent raters conducting the evaluation and a third resolving any discrepancies.

### Analysis

All statistical analyses were performed using RStudio 2021.09.1, using two-tailed tests for significance, if no a priori hypotheses indicated the use of one-tailed tests.

*Dream rating.* Audio recordings of the dream recall and retrieval of the audiobook content were transcribed for the analysis. Three independent blind raters were asked to match the written dream reports with one of the four audiobook conditions. They were blind to the audiobook condition and all subject details, uninvolved in data collection or audio transcription, but had thoroughly studied the content of each audiobook, having both listened to them and read the text versions in detail (which were adapted to match the exact content of the audiobooks). Therefore, they had a detailed knowledge of the four narratives. While performing their ratings, they had access to both the audiobooks and the text versions, with timestamps for each page to facilitate comparisons between the text and the audiobook timing. Additionally, they were provided with information for how long participants had listened to the audiobook, enabling them to compare this with the timestamps in the text version. To avoid potential bias, for the rating they were presented with single dream reports in a randomized order. More specifically, dream reports were randomized across all participants, such that reports from the same participant were distributed throughout the entire dataset and interspersed with those of other participants. The randomization order was unique for each rater. Each report included the transcribed version of the free dream recall and additional information about the content of the dream collected through a standardized cued dream questionnaire. Information in the transcribed audio recording that could indicate that dream reports belonged to the same participant or revealed the participant’s identity was removed beforehand (e.g. “like in my previous dream about…”). For their rating, they could thus only take into account information from one report at a time. The raters were asked to rate the confidence of their decision according to how much information about the audiobook they detected in the dream report (0 = guessed, 1 = gut feeling, 2 = implicit indication, 3 = explicit indication). If raters stated to have traced an implicit or explicit indication for the listened audiobook, they were asked to report which part of the audiobook narrative was incorporated. Please note that not all narratives of dreams for which ratings indicated a direct or an indirect incorporation could be linked to specific events in the audiobooks. For a full list of dreams with indirect and direct incorporations that could be traced to specific events in the audiobooks, see Supplemental Material (Table S7a-c). These dreams represent all direct and indirect incorporations for which all three raters agreed there was an incorporation. To assess whether the information that is reprocessed during dreams indeed stems from the most recently listened audiobook passage, we examined the temporal origin of the explicit and implicit references to audiobook content that could be traced to specific sections of the audiobook (Table S7b-c).

*Concordance analyses.* The agreement between rated and the actual audiobook was assessed across the whole night by using Cohen’s kappa (concordance measure between the rated condition and the ground truth). If there is significant concordance between the rated and the actual conditions, information about the learning material must be present in the dream reports. Concordance measures are expected to be low in these analyses due to anticipated variability in memory incorporation across dream experiences, possible inaccuracies in the raters’ recall of narratives, and differences in their criterion for identifying distorted forms of the audiobook narrative in the dream reports. Two participants listened to two audiobooks throughout the night because of long wake times. In these cases, ratings were considered correct when the rated audiobook had been presented prior to any previous awaking during the night.

*Individual ratings*. In a first analysis, each decision made by the three blind raters for every dream report was considered individually (a total of 182 ratings for the 61 obtained dream reports). The inter-rater agreement was positive and significant, indicating they based their ratings on similar information in the dream reports (ĸ = 0.236, z = 5.48, p < 0.001). Please note that given the noise sources stated above, also inter-rater agreement is expected to be low in this analysis.

*Combined rating*. In a second analysis, the ratings for each dream report were combined across raters to obtain one final decision on which audiobook had been previously listened to. For this, the subjective confidence of ratings was taken into account. For each dream report, confidence scores were summed for identical ratings across raters. The audiobook category reaching the highest total confidence score was selected as the consensus rating for that dream report. This process resulted in 43 consensus ratings for the 61 obtained dream reports. If two books had the same total confidence, no consensus was reached, and the report was treated as missing value.

*Effects of sleep stage and time of night.* The same analyses were then repeated after dividing the dataset by the last sleep stage experienced prior to awakening (NREM and REM sleep) and after dividing the dataset by early and late sleep awakenings. Sleep periods in which participants were woken up from S2 or SWS were considered NREM awakenings, and sleep periods when participants were awoken from REM sleep REM awakenings. The first two awakenings of the experimental night were considered early awakenings while the last two awakenings of the night for each participant were considered late awakenings. Thus, when a participant was woken up five times, the third awakening was not considered in the early vs. late awakenings analyses. Contingency tables were generated to visualize results for all analyses.

*Per participant rating.* In an additional, consecutive but separate round of dream ratings, the raters were presented with the collected dream reports of all individual participants, de-blinding which dream reports belonged to the same participant (and thus the same audiobook condition). Again, they had to rate which audiobook a person had listened to throughout the night.

*Incorporation strength.* We calculated a measure for the amount of information about the previously listened audiobook represented in later dream content. For each dream report, we summed the confidence ratings provided by the three raters which, based on the instructions, reflect the extent of information related to the audiobook content. Confidence statements for correct decisions were entered as positive values, statements for incorrect decisions were entered as negative values into this reprocessed information score.

The Wilcoxon signed rank test was used to test potential differences in the amount of dream incorporations between NREM and REM, as well as early and late sleep awakenings. NREM awakenings included all the awakenings in which participants woken up from S2 and SWS. For this analysis, only participants with dream reports from both NREM and REM awakenings were considered, resulting in a reduced dataset with dream report-related data paired by sleep stage for each participant (remaining N = 7). The data from the first two and last two awakenings of the night were averaged for each participant to obtain a final information score for early and late night. For this purpose, participants without any awakening in the first or second part of the night were excluded from this analysis (remaining N = 12). Given the low sample size, these comparisons should be interpreted with caution.

*Correlations with memory performance.* Dream incorporation of previous learning material has been associated with better memory for the content in previous studies^27,29^. We thus tested whether dream incorporation is *positively* related with memory for the audiobook. We used a non-parametric correlation coefficient that is suited for small samples (Kendall’s tau, one-tailed test for τ > 0) to correlate memory performance for the listened audiobook passage, measured as percent correctly recognized audiobook content, with the amount of information about the book reprocessed during dreams in that sleep period. For this analysis, only observations with a positive information score were included, averaged at the participant level (N = 11). The same test was then repeated for the free recall performance.

We then used a linear regression model (*lm* function) to explore whether the occurrence of NREM or REM sleep before each awakening could explain the association between dream incorporation and memory performance. In our model, audiobook recognition was entered as the dependent variable, while information score and sleep stage were entered as possibly interacting predictors (model used: *lm(audiobook_recognition ∼info_score*last_sleep_stage*). The variance inflation factor (VIF) analysis yielded a score of 3.95 for the variable last sleep stage and 2.20 for info score, indicating no severe collinearity issues that would justify excluding the model. Following the same rationale, we tested whether early or late night could predict the memory benefit for instances with stronger incorporations of the audiobook (model used: *lm(audiobook_recognition ∼info_score*night_segment*). The VIFs for *night segment* and *info score* were 4.06 and 5.38, indicating moderate collinearity. Given the exploratory nature of the analysis, the model was nevertheless included. To retain as many data points as possible, for these analyses we kept all dreams with positive information score as separate observations (N_sleepstages_ = 12, N_nightsegment_ = 14).

Given the low sample size, these correlations and results of the linear regression models should be interpreted with caution.

Furthermore, we controlled whether the presence of both NREM and REM sleep prior to awakening influenced dream incorporation and memory performance. First, we categorized awakenings based on whether they followed only NREM sleep or both NREM and REM sleep. We then aggregated data at the participant level, and used Wilcoxon signed rank tests to compare the *information score*, as well as the *audiobook recognition* and the *free recall* measures between awakenings with the occurrence of both NREM and REM sleep, or only one of these sleep stages. For this purpose, participants who did not experience both conditions were excluded from the analysis (paired data: N_infoscore_ = 9, N_recognition_= 8, N_recall_= 8).

Finally, Kendall’s Tau was used to test whether the great variability in audiobook listening time across participants influenced dream incorporation strength.

*Laboratory setting incorporation.* For each dream report, we also evaluated whether we could detect any incorporation that was related to the experimental situation (experimenter, EEG cap, experimental beds, experimental procedure, etc.). Two separate raters independently evaluated the presence of experimental incorporation into dreams (1 = *expsit yes*, 0 = *expsit no*). In case of disagreement between the two raters, a final consensus was reached through discussion, following an approach previously employed, when quantifying dream incorporations^28^. An example of absent, implicit and explicit laboratory dream incorporation is provided in the Supplemental Material section (table S9). Note that differently from the audiobook-related dream rating where raters provided confidence scores, this categorization is purely qualitative, and provided for descriptive purposes only.

We first used a chi-square test to examine the frequency of both incorporation dimensions across all dreams (N = 61). For this purpose, each dream report was treated as a single observation. We then computed a Wilcoxon rank sum test to compare the amount of audiobook-related information (*information score*) reprocessed in dreams with and without *expsit* incorporation (N = 61). In this case, Wilcoxon rank sum test was chosen instead of Wilcoxon signed rank, as using a paired version of the dataset with aggregated data points would have resulted in substantial data loss, leading to only 5 paired observations. For each participant, we then calculated the overall number of dreams with experimental situation incorporation and dreams with correct audiobook identification, and used Kendall’s Tau to test the relationship between the two measures (N = 19). We additionally correlated, for each participant, the overall ratio of dreams showing *expsit* incorporation over dreams without any, with the overall ratio of dreams showing audiobook incorporation divided by the number of dreams where no audiobook incorporation was reported (N = 19). Since it remains unclear whether the incorporation of the experimental setting interferes with or facilitates content incorporation [75,79], we conducted two-tailed tests for these analyses. Finally, we wanted to examine the temporal occurrence of such incorporations across the night and sleep stages. The McNemar test was used to compare the frequency distribution of dreams with and without *expsit* between early and late sleep (N = 12), as well as between NREM and REM sleep awakenings (N = 7). Following the same rationale used for the comparison of information scores across the night, only participants who had dream reports from both night segments or from both NREM and REM awakenings, respectively, were included. This resulted in two reduced, paired datasets. Given the low sample size, the results of these analyses should be interpreted with caution.

*Dream length and audiobook incorporation.* To test whether the length of dream reports determined the amount of audiobook-related information blind raters detected in dreams, we calculated Kendall’s Tau. Dream length was quantified as number of words in each dream report. Dream reports were redacted such that they only included content referring to the dream experience. For this analysis, data was averaged across participants (N = 11).

## RESULTS

### Descriptives

Each participant was awoken multiple times throughout the night (4.1 ± 0.44 [mean ± SD], N = 20). Out of 82 total awakenings (N_NREM_ = 56, N_REM_ = 10, N_S1_ = 5, N_wake_ = 11), 61 dream reports were collected (N_NREM_ = 35, N_REM_ = 11, N_S1_ = 5, N_wake_ = 10). For reports from NREM awakenings, 32 were collected from S2 and 3 from SWS. On average, participants had 3.1 ± 1.7 [mean ± SD] of total dreams during the night. On some occasions, participants reported more than one dream after being awoken. This occurred for 9 awakenings in total, during which participants would report 2 or a maximum of 3 dreams at once, resulting in a total of 11 additional dream reports. Multiple dreams reported upon a single awakening reflected distinct storylines and were perceived by participants as temporally separate experiences. Accordingly, separate responses were recorded in the dream questionnaire. In total, dream content was reported in 50 awakenings, making the actual dream report rate 61%. One participant had no dreams to report throughout the night and was thus excluded from all analyses (remaining N = 19).

During an average sleep period (across all awakenings), they were awake for 7 ± 2 minutes, spent 8 ± 3 minutes in sleep stage 1 (S1), 29 ± 12 minutes in sleep stage 2 (S2), 14 ± 7 minutes in slow wave sleep (SWS), and 16 ± 9 minutes in REM sleep, giving an average total sleep time of 67 ± 13 minutes [mean ± SD]. For additional details, see Supplemental Material (fig. S1, tables S1-S6).

### Audiobook-related memory is incorporated into dreams

To assess whether the specific content of the audiobook encoded before sleep is reprocessed during subsequent dreams, three independent blind raters were asked to judge which audiobook each participant had listened to before experiencing a dream, based on the transcript of the dream report and their answers to the subsequent structured dream questionnaire (see Methods). For each dream assignment, the raters also indicated their confidence in the decision using a four-point scale (0 = guessing, 1 = gut feeling, 2 = implicit indication, 3 = explicit indication). We initially considered each decision made by the three blind raters for every dream report individually. This approach was chosen to mitigate potential data loss due to ambiguous ratings when aggregating the individual raters’ decisions. We compared these individual ratings to the actual experimental conditions (ground truth) and tested for the concordance of rater decisions. When considering the individual rating and comparing it to the actual experimental condition, dream reports could be successfully matched with the corresponding audiobook condition (individual ratings: ĸ = 0.13, z = 3.07, p = 0.002; Fig 1C). We then calculated a combined rating of all individual raters for each dream report, considering both the frequency of choice and the confidence of their rating to confirm the obtained result (see Methods). Again, we observed a significant concordance of combined ratings and the actual audiobook condition (combined rating: ĸ = 0.2, z = 2.37, p = 0.02; Fig 1E). The raters could decide which audiobook a person had listened to before falling asleep, based solely on the content of the dream report.

**Figure 1.**
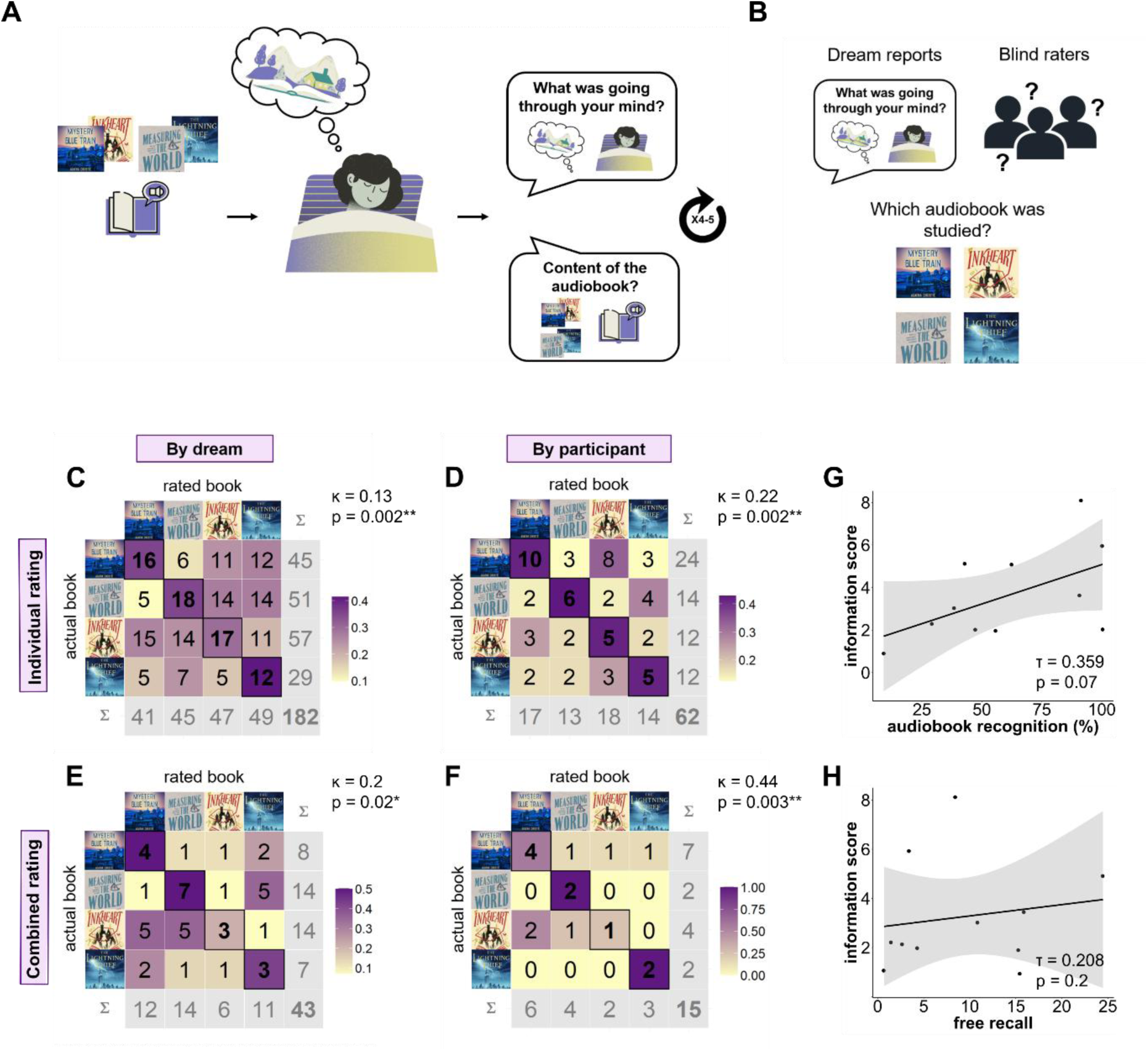
Incorporation of audiobook-related information into dreaming A. Experimental design. Participants listened to one out of four audiobooks before falling asleep. Every 90 minutes they were woken up and asked to report their dreams. They were also tested for how well they could remember the previously encoded audiobook passage. The procedure was repeated across the night up to 4-5 times. **B. Method.** Three blind raters evaluated the dream reports. For each dream they were asked to guess which audiobook participants had studied, based solely on the dream reports, and to indicate their confidence, based on how much audiobook-related information they detected. **C.** Confusion matrix of listened vs. rated audiobook for individual ratings by dream. Total number of cases: 182, 1 missing value (missing rating). **D.** Confusion matrix of listened vs. rated audiobook for combined ratings by dream. Total number of cases: 43, 18 missing values (ambiguous ratings, no concordance found between the three raters). In all confusion matrices, the color scaling is determined by weighting each case according to the total number of actual instances within each audiobook condition. Note number of correct rater decisions on the diagonal. **E.** Confusion matrix of listened vs. rated audiobook for individual ratings by participant. Total number of cases: 62, 1 missing value (missing rating). **F.** Confusion matrix of listened vs. rated audiobook for combined ratings by participant. Total number of cases: 15, 5 missing values (ambiguous ratings, no concordance found between the three raters). **G-H.** Relationship (Kendall’s tau) between the amount of information reprocessed in dreams (information score) and the memory strength for the listened to audiobook passage. The information score represents the sum of the adjusted confidence of the three blind raters for each rated dream. Only positive scores--indicating a cumulatively correct audiobook assignment--were included in this analysis (for further details, see the Analysis paragraph “*Incorporation strength*”, included in the Methods section). G: audiobook recognition in % recognized, the asterisk depicts a significant correlation, one-tailed test. H: free recall in number of words per minute of audiobook listened. The relationship was not significant in a one-tailed test). * p < 0.05, ** p < 0.01

We then provided the blind raters with all dream reports belonging to the same participant, instructing them to repeat their rating, this time for each dream set, thus making only one rating of audiobook condition per participant. The raters were aware that each participant listened to the same audiobook throughout the night, making it possible to infer a participants’ experimental condition from a single incorporation of audiobook content across all the dream reports. As expected, the raters were well able to assign the correct audiobook condition for the participants (individual ratings: ĸ = 0.22, z = 3.07, p = 0.002; combined rating: ĸ = 0.44, z = 3.0, p = 0.003; Fig. 1D-F).

Finally, we split the data according to the last sleep stage participants were in prior to each awakening to investigate dream incorporation from NREM and REM sleep awakenings separately. Most of the dream reports came directly after NREM sleep (n=35), while 11 dreams were reported after REM sleep (see Table S4 in Supplemental Material). This distribution accurately reflects the distribution of sleep stages across the night. In the non-aggregated dataset, dream reports after both NREM and REM sleep reliably allowed raters to predict which audiobook had been listened to before (individual ratings: NREM sleep: ĸ = 0.13, z = 2.39, p = 0.02; REM sleep: ĸ = 0.312, z = 2.96, p = 0.003; Fig. 2A-B). This was confirmed for REM sleep awakenings in the combined rating (combined rating: REM sleep: ĸ = 0.41, z = 2.52, p = 0.01; NREM sleep: ĸ = 0.17, z = 1.43, p = 0.15; see Fig S2A).

**Figure 2.**
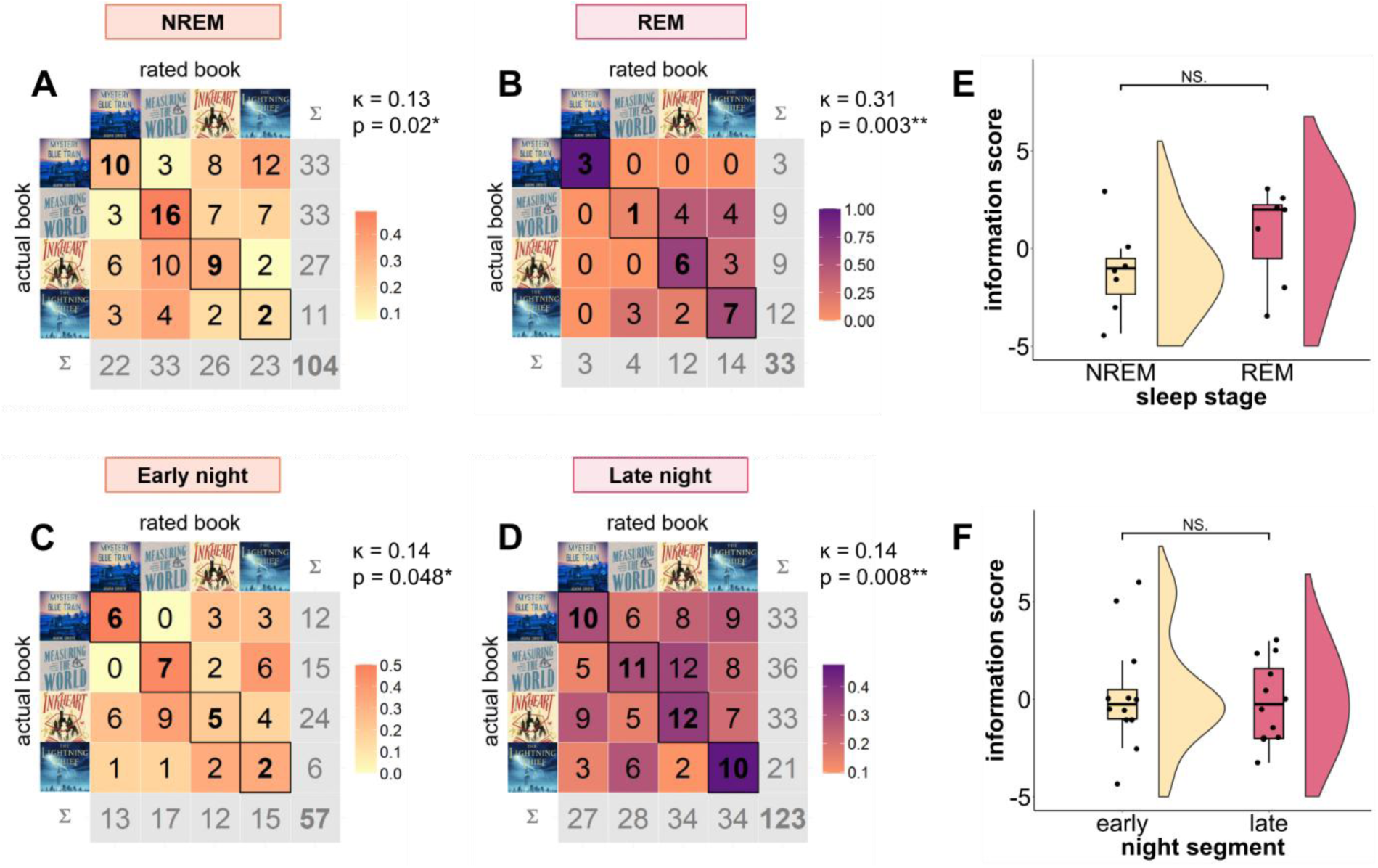
**Incorporation of audiobook-related information into dreaming, night dynamics A-B. NREM and REM sleep awakenings**. Total number of cases: 104 and 33, 1 missing value in NREM (missing rating)**. C-D. Early and late awakenings**. Total number of cases: 57 and 123. In all confusion matrices, the color scaling is determined by weighting each case according to the total number of actual instances within each audiobook condition. Note number of correct decisions on the diagonal. In all confusion matrices, the color scaling is determined by weighting each case according to the total number of actual instances within each audiobook condition. Note number of correct rater decisions on the diagonal. **E.** Amount of audiobook-related information reprocessed in dreams by NREM and REM sleep awakenings. The difference is not significant (N.S). **F.** Amount of audiobook-related information reprocessed in dreams by early and late sleep awakenings. The difference is not significant (N.S). In the two plots depicting group comparisons, the horizontal black lines within the boxplots represent the median value, whereas the dots represent the single observations. * p < 0.05, ** p < 0.01

As the time of night, beyond sleep stages, shapes which memory sources contribute to dreaming [67], we then split the data into dreams that occurred during the early night and the late night. It was possible to correctly identify the encoded audiobook based on dream reports from both the early and late night when considering each rating separately (individual ratings: early night – first two awakenings: ĸ = 0.14, z = 1.98, p = 0.048; late night – last two awakenings: ĸ = 0.14, z = 2.66, p = 0.008; fig 2C-D). Again, this was confirmed in the combined rating, yet only for reports collected in the late night, when REM sleep is predominant (combined rating: early night – first two awakenings: ĸ = 0.18, z = 1.4, p = 0.16; late night – last two awakenings: ĸ = 0.24, z = 2.22, p = 0.03; see Fig. S2B). The discrepancy in results for NREM and early sleep when considering the combined ratings may be due to consistent data loss caused by unresolvable ambiguous ratings. Since the signal was weaker for NREM sleep and the early night, this ambiguity may have rendered the effect undetectable in the combined dataset.

To better quantify the amount of audiobook-related information represented in dreams, an information score was obtained for each dream by weighting the confidence of the three raters by the correctness/incorrectness of their decision (see Methods). This served as a quantitative measure of reprocessing in the dream, indirectly reflecting how close the dream content was to the episodic narrative in the audiobook passage. The total audiobook listening time did not influence how much audiobook information participants later incorporated into dreaming (τ = 0.09, z = 0.43, p = 0.66, N = 13).

We compared the amount of information about the previous learning episode available in each dream report between NREM vs REM, and between early sleep vs late sleep awakenings. Statistically, there was no significant difference between NREM and REM sleep (Z = 8, p = 0.31, CI [-5.67, 2.50], N_paired_ = 7, Fig. 2E, note that participants with missing data in one of the brackets were excluded), nor between early sleep and late sleep awakenings (Z = 41.5, p = 0.84, CI [-1.67, 2.83], N_paired_ = 12, Fig. 2F, note that participants with missing data in one of the brackets were excluded). These findings suggest that previous learning episodes resurface in a comparable fashion during conscious experience during both NREM and REM sleep throughout the whole night. However, when looking at NREM and REM sleep separately in early and late sleep, only dream reports from late sleep could be reliably assigned by the raters (late night - NREM: ĸ = 0.183, z = 2.58, p = 0.001; REM: ĸ = 0.3, z = 2.52, p = 0.01; early night - NREM: ĸ = 0.101, z = 1.15, p = 0.245; REM: ĸ = 0.333, z = 1.53, p = 0.13; fig. S3A, all results pertain to the individual ratings, see Fig. S3B for the combined rating). Taken together with the above-mentioned concordance analyses and incorporation findings, this suggests that while the strength of incorporated information may not vary throughout the night, incorporations might occur more frequently in dreams experienced during the late night.

### Specific audiobook-related content can be detected in dream reports

In seven cases (representing 11% of all dream reports), all three raters agreed on detecting indirect or direct incorporations of the audiobook content (see Table S7a-c). In all of these cases, the information could be traced back to the passage of the audiobook that was listened to during the waking time immediately before the dream occurred. In two of these cases, more remote events were additionally incorporated.

### Correlating dream incorporation and memory for the audiobook

If dreaming reflects memory reprocessing, dreaming of the audiobook should improve memory for its content. To test this, we analyzed whether the amount of information about the previous learning episode available in dream reports was related to better memory for the corresponding audiobook passage. For each participant, we thus correlated the information score, as a measure for memory reactivation, with how much of the previously encoded audiobook passage they recognized in a later retrieval test. This analysis included only instances for which the correct audiobook had been correctly assigned by the raters, averaged per participant across the whole night. Although not statistically significant, we observed a tendency for the amount of information reprocessed during dreams to correlate with later recognition memory for the listened audiobook passage (τ = 0.359, z = 1.5, p = 0.07 one-tailed, N = 11; Fig. 1G). Reprocessing of previous learning material through dreaming activity thus tended to be positively associated with audiobook recognition, which suggests an association between dreaming and memory consolidation. The same effect did not emerge for free recall performance (τ = 0.208, z = 0.87, p = 0.2 one-tailed, N = 11; Fig. 1H). The occurrence of NREM or REM sleep before each awakening, as well as the distinction between early or late night dreams, could not explain the relationship between audiobook recognition and information score (Regression models, 1) NREM-REM: F(3, 8) = 5.02, p = 0.03, R² = 0.65, Adjusted R² = 0.52, interaction *info score*-*last sleep stage*: beta =-0.09, 95% CI [-0.23, 0.04], t(8) =-1.57, p = 0.155; 2) Night segment: F(3, 10) = 2.30, p = 0.14, R² = 0.41, Adjusted R² = 0.23, interaction *info score*-*night segment*: beta = 0.006, 95% CI [-0.20, 0.21], t(10) = 0.07, p = 0.94. See Table S8a-b for additional information).

Neither dream incorporation nor the memory measures differed significantly between awakenings after experiencing both NREM and REM sleep, or only one of these sleep stages (*information score*: Z = 31.5, p = 0.28, CI[-1, 2.3], N_paired_ = 9; *audiobook recognition*: Z = 15, p = 0.67, CI[-0.4, 0.32], N_paired_ = 8; *free recall:* Z = 24, p = 0.4, CI[-8.8, 4.9] N_paired_ = 8).

### The experimental situation gets incorporated into dreams

Two additional blind raters were asked to evaluate whether they could observe any incorporation of the laboratory environment or the experimental setting in the dreams. Contrary to our analysis on audiobook incorporation, no ground truth can be provided for incorporation of the experimental situation into dreaming, because all participants underwent the same experimental procedure. We can thus not draw statistically verifiable conclusions on whether or not an incorporation of the experimental situation actually occurred, a problem that also past research on dream incorporation has faced [75]. In a total of 61 dream reports, the blind raters identified at least one element of the experimental situation in 20 dreams, while remaining 41 showed no evidence of experiment incorporation. Dreams with incorporation of the experimental situation thus represented 33% of the total pool, which conforms to incorporation rates reported in the literature [75] (Fig. 3A; see Fig. S5-S6, and Table S9 for further details).

**Figure 3.**
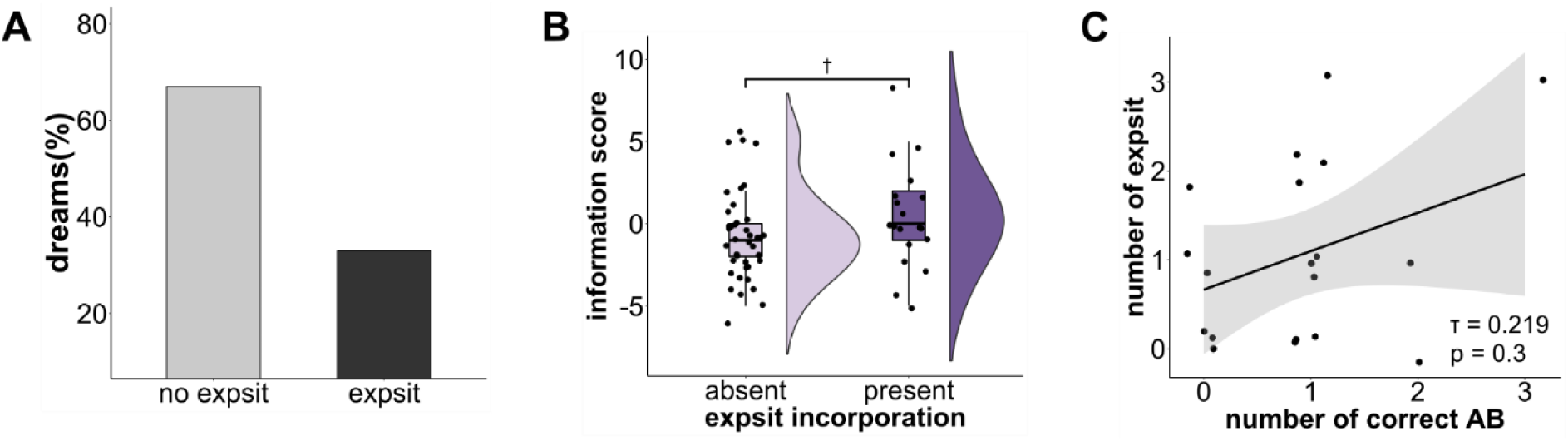
Incorporation of the experimental situation into dreaming. **A.** Proportion of dreams with and without incorporation of the experimental situation (*expsit*). **B.** Amount of audiobook-related content (information score) in dreams with and without *expsit* incorporation. **C.** Relationship between the number of detected audiobook incorporations (AB) and *expsit* incorporations by participant (Kendall’s tau). In the plot depicting group comparison, the horizontal black lines within the boxplots represent the median value, whereas the dots represent the single observations. Crosses indicate a trend in the group comparison (p < 0.1).

When looking at the relationship between experimental incorporation and reprocessing of audiobook-related content, dream reports with experiment incorporation did not include audiobook incorporation more frequently than dreams with no presence of the laboratory setting (χ²(1, N = 61) = 1.37, p = 0.24). However, we observed a tendency in dreams reactivating the experimental situation to incorporate more audiobook content, reflected by a comparatively higher *information score* than in dreams showing no incorporation of the experimental situation (U = 529.5, p = 0.06, CI [-0.001, 3.0], N = 61, Fig 3B). At the participant level, no significant association between the frequency of experimental situation incorporation and audiobook incorporations was found (absolute numbers of incorporations: τ = 0.219, z = 1.06, p = 0.3 two-tailed, N = 19; Fig. 3C. Ratio, corrected by instances without incorporations: τ = 0.16, z = 0.82, p = 0.4 two-tailed, N = 19).

## DISCUSSION

### Dream incorporation of the learning material and memory reprocessing

We find that blind raters can correctly assign which audiobook someone had listened to before falling asleep, based solely on the dreams they reported during the night. The content of the audiobook thus resurfaced in dream narratives during consolidated sleep. This dream incorporation may reflect spontaneous neural reactivation of learning-related activity patterns [18–22]. When assessing audiobook reprocessing in the current dataset at the neural level, we observed that brain activity during REM sleep contained information about the previously learned audiobook and was associated with the retention of the audiobook content [results reported in a separate manuscript, 86]. Moreover, participants who incorporated the audiobook during the night showed a stronger neural reinstatement. Taken together, these findings support the idea that offline neural reprocessing surfaces in conscious experience during sleep, thereby contributing to dream construction.

We found that participants who incorporated more audiobook content into their dreams also tended to better recognize the audiobook passages they studied before. Findings regarding the association between dream incorporation and memory performance have been mixed (see [28] for an overview). In our study, the effect of dream incorporation on memory performance at the participant level was weak and did not reach significance in the small sample, nor was it consistent across memory measures. The indirect nature of how memory reactivation surfaces in the content of dream experiences and the resulting uncertainty in quantifying the amount of reactivation, as well as the challenge of assessing memory performance for naturalistic stimulus material may have additionally diluted the effect. Our results are in line with previous studies that reported a positive effect of dream incorporation on memory retention for two different kinds of learning material, word-picture learning and spatial navigation performance [27–29], yet highlight the difficulty of detecting a clear association between dream incorporation and memory performance. Our findings indicate that also episodic narratives, content similar to our continuous waking experience, resurface in conscious experience during sleep and that this incorporation may be related to the memory for these stories. Studies with larger sample sizes and experimental designs tailored to assess the effect of dream incorporation on memory performance are needed to corroborate this result.

### A novel paradigm to study dreaming

For this study, we took a novel approach to study memory incorporation into dreaming. Similar to Schoch et al. [27], we explicitly manipulated the content of learning material to bias memory processing during sleep, and consequently in dreaming. We extend that work by showing that also complex naturalistic stimuli, i.e., the narratives of audiobooks, become incorporated into later dreams. Contrary to previous studies, where participants were asked to retrospectively rate the similarity of their dream reports with their daily activities in the days or weeks preceding the experiment [40,41,72,87], more and more recent studies employ independent raters who are instructed to judge the reported dream contents [26–29,88]. Our study employed three independent and blind raters to make alternative forced-choice judgements about the content of the previous learning material, allowing us to verify their ratings against a ground truth. This makes it possible to directly test the hypothesis whether learning content is incorporated into dreams during the night and investigate conscious experience during sleep as a marker of memory reactivation.

Incorporation rates in studies employing independent raters are lower, and are necessarily more conservative than incorporation rates in studies where participants rate concordance of waking life and dreams. Contrary to other approaches, alternative forced choice judgements create noise in the data in cases where no memory incorporation occurred in dreaming and raters have to guess conditions. It is therefore remarkable that we were able to detect a memory incorporation signal in dream reports at a level that exceeds chance.

### Characteristics of dreams in different sleep stages and at different times of the night

Learning-related information was reflected in the content of dream reports originating from both NREM and REM awakenings, without appreciable differences between early and late night. This seems to contrast previous observations ascribing incorporations of recent memories and waking-life experiences mostly to the early night [66–68], and findings from reactivation of learning-related neural activity in animals that report that the strongest reactivation of learning activity occurs during the first 30 min of (NREM) sleep [89]. Our multiple awakening paradigm entailed sequential encoding of the audiobook throughout the night, thus ensuring temporal proximity to the originating wake experience during all times of the night. In all cases when our blind raters detected an audiobook-related incorporation, the reactivated content could be traced back to events in the audiobook section presented immediately before the sleep period in which the dream occurred. Moreover, the amount of incorporated information did not differ between sleep stage or time of the night. This indicates that temporal proximity to the encoded experience may be a more decisive factor in whether information gets incorporated into dreams. However, when analyzing incorporations from NREM and REM awakenings during early and late sleep, only late-sleep dream reports were reliably linked to the learning condition. This suggests that while the amount of incorporated information remains consistent, incorporations or detectable audiobook-related content may be more frequent in late-night dreams. More frequent incorporations late during the night might be attributable more to the ability to recall dream content than to an influence on dream construction itself. In line with this reasoning, we found a positive correlation between dream length and information score (see Supplemental Material, fig. S7).

Note that in our study, we did not characterize the memory sources of the dream reports (e.g., autobiographical, semantic) through systematic coding, thus making it impossible to distinguish these qualitative features of dreams and compare them between NREM and REM sleep awakenings. Instead, our method relied on forced choice decisions made by blind raters who indicated what sources of information they based their rating on (guess, gut feeling, indirect or direct incorporation), and thus in which way content from the audiobook was incorporated into dreaming. Only 2% of the dreams contained an explicit incorporation of the audiobook-content, in line with previous evidence reporting episodic incorporation into dreaming only 2-4% of the time [40–42]. In 10% of the cases, raters could instead detect indirect references in the dream report, which could be traced back to specific occurrences or themes in the book. While explicit dream references refer to unambiguous episodic events that occurred in the book narrative, implicit references encompass a broader spectrum of mixed memory sources, including semantic similarity, as well as related autobiographical sources. Our findings align with other studies that made no distinction in incorporation type [27], which similarly found no significant difference in incorporation rate between NREM and REM awakenings. Furthermore, it is not possible to determine when dreaming exactly occurred prior to each awakening. Our analyses were performed and results are interpreted under the assumption that all dreams occurred during the few minutes before waking up. For this reason, our findings comparing dream reports from NREM and REM awakenings do not rule out the possibility that memory incorporation leaked into NREM or REM dream awakenings from preceding sleep stages. More generally, when no stage suppression or manipulation is operated by experimental design [58], results on the effects of sleep stages on dreaming need to be interpreted with caution.

### Dream incorporations of the experimental situation

In our study, approximately 30% of the dream reports contained incorporations of the experimental situation. This is in line with the dream incorporation frequency usually reported, ranging between 30-70% when both explicit and implicit elements of the experimental situation are considered [75].

Spending the night in a sleep laboratory and performing an experimental task is a salient, novel experience, which may lead to competition between episodic features of this personal event and the audiobook-related content for neural resources during night time reprocessing [90,91]. When assessing the distribution of audiobook and laboratory setting incorporations across all dream reports, we did not observe any significant pattern. This suggests that no substantial interference between the two types of information occurred in dreaming. However, we observed that dreams reporting at least one element of the experimental situation participants underwent also tended to contain a higher amount of audiobook-related information. To assess whether this trend was driven by some participants, who may be overall ‘stronger incorporators’ of waking-life elements [36], we examined the association between number of audiobook-and experiment-related incorporation at the participant level, yet found no significant relationship. While our findings do not allow definitive conclusions about the relation between laboratory and stimulus-related dream incorporations, we do not find strong evidence for competition. Further research is needed to investigate this connection more closely.

Since the laboratory setting provides the broader context for studying the learning material, its incorporation into dreams might reflect the general reactivation of the pre-sleep experience. Examining the interplay of learning content and experimental setting incorporation across different kinds of task, particularly considering how relevant the experimental context is to task completion, may further elucidate the nature of this interaction.

### Limitations

It is important to note that due to the overall small sample size, some of our analyses may be underpowered. Effects that only emerged subtly in our study should be further examined in larger samples, e.g., the relationship between dream incorporation and memory. Additionally, the small sample size restricted our ability to account for inter-individual differences, which could have influenced the dream incorporation dynamics observed across the night. With a larger sample, mixed-effect models would be more suited to control for such random effects in the data.

Furthermore, in the multiple awakening paradigm used in this study, participants underwent encoding multiple times throughout the night, immediately before falling asleep. This method does not provide a baseline measure of participants’ initial memory strength before sleep, making it more difficult to determine who benefitted from audiobook-related dream incorporation.

However, this approach allowed us to explore whether the temporal proximity between learning and dreaming influences which content is more likely to be incorporated into dreams. Except for two instances, for explicitly and implicitly incorporated information, the dream content only referred to the most recently listened portion of the audiobook. Note, however, that it was not possible to precisely link other incorporations (i.e., correct ratings judged with “gut feeling”) to specific audiobook passages. As a result, these incorporations may not be uniquely attributable to the most recent audiobook segment: both the retrieval of earlier-encoded content and the encoding of new material during each awakening may have shaped subsequent dream narratives. Overall, this limits our ability to determine the actual contribution of temporal proximity in learning-related incorporation across the full dream set.

Moreover, given the current experimental design it is not possible to fully rule out that the final minutes of audiobook encoding occurred as participants were already drifting off. If this were the case, some instances of content incorporation into subsequent dreams might depend on active dream incorporation, not memory reactivation, which cannot be disentangled in this study.

Another potential methodological limitation lies in the dream recall and free recall procedures, subsequent processing of free speech records, and resulting word count measures. To ensure standardization across all transcribed texts, with regards to speaking style, verbosity, and language use, we applied a stepwise approach to redact the original transcripts, following common guidelines to enhance comparability across participants’ recounts. However, operator-dependent biases on resulting word count measures that constituted estimates for dream length and free recall performance cannot be entirely ruled out. In future studies, the use of automated methods for linguistic processing, in support of manual operations, could be considered to help reduce these potential biases.

Relatedly, our audiobook recognition score only provides a coarse measure of memory performance. Due to the nature of the experimental paradigm, we did not assess participants’ memory sensitivity for the audiobook passages, but report hit rates for recognition performance. Although freely recalled information and audiobook recognition positively correlated (see fig. S8), we cannot entirely rule out the possibility that participants who incorporated audiobook material more strongly would also have mistakenly recognized additional audiobook passages due to response biases. In light of these considerations, the observed findings on dream incorporation and memory consolidation should be interpreted with caution.

Finally, relying on dream reports currently remains the primary available method for accessing the informational content of the dream experience itself. As a result, at present it is not possible to disentangle the memory benefit of dream incorporation per se from the benefit of retrieving incorporated dream content.

## CONCLUSIONS

Our results indicate that dreams reflect internal memory reactivation processes. Both dreams from NREM and REM sleep, as well as from the early and late night contained information about studied audiobook narratives, indicating that memory incorporation is a universal process, occurring during all sleep stages and regardless of when sleep occurs. The more information about the audiobook could be detected in dreams, the better participants tended to recognize the narrative across the sleep period. Our experiment was designed to mirror paradigms investigating neural reactivation in humans, making it possible to relate findings on dream incorporation and neural reactivation [21,22,86,92]. Indeed, a paper assessing neural reactivation in our paradigm was able to detect reprocessing of audiobook content in our participants’ electrical brain activity during sleep [86]. Intriguingly, neural reactivation was stronger in participants who dreamt of the audiobook, lending further strength to the idea that dreaming at least in part reflects nighttime memory processing. While it is becoming more and more apparent that memory reprocessing, consolidation, and dreaming are strongly intertwined, it is still unclear how they interact, what determines which memories will resurface in dreaming, or which type of processing these fragmented memories undergo. With recent methodological advances, future research incorporating larger sample sizes and preregistered hypotheses may soon be able to address these gaps [92].

## ACKNOWLEDGMENTS

We would like to thank Dr. Philipp Paulus for assistance with illustrations.

## AUTHORS’ CONTRIBUTIONS

Conceptualization: SG, MS

Methodology: JP, SG, MS

Investigation: SKA, MS, SG, MS

Visualization: JP

Funding acquisition: SG, MS

Formal Analysis: JP, SJW, DK, MS

Project administration: MS, SG

Supervision: MS

Writing – original draft: JP, MS

Writing – review & editing: JP, DK, SG, MS

## DATA AVAILABILITY STATEMENT

Part of the data collected in this project is available in the DREAM database (Wong et al., 2023), https://doi.org/10.26180/22133105.v5. Behavioral data and codes are available at the Open Science Framework repository (https://doi.org/10.17605/OSF.IO/EM5AC).

## Supplemental Material

**Figure S1.**
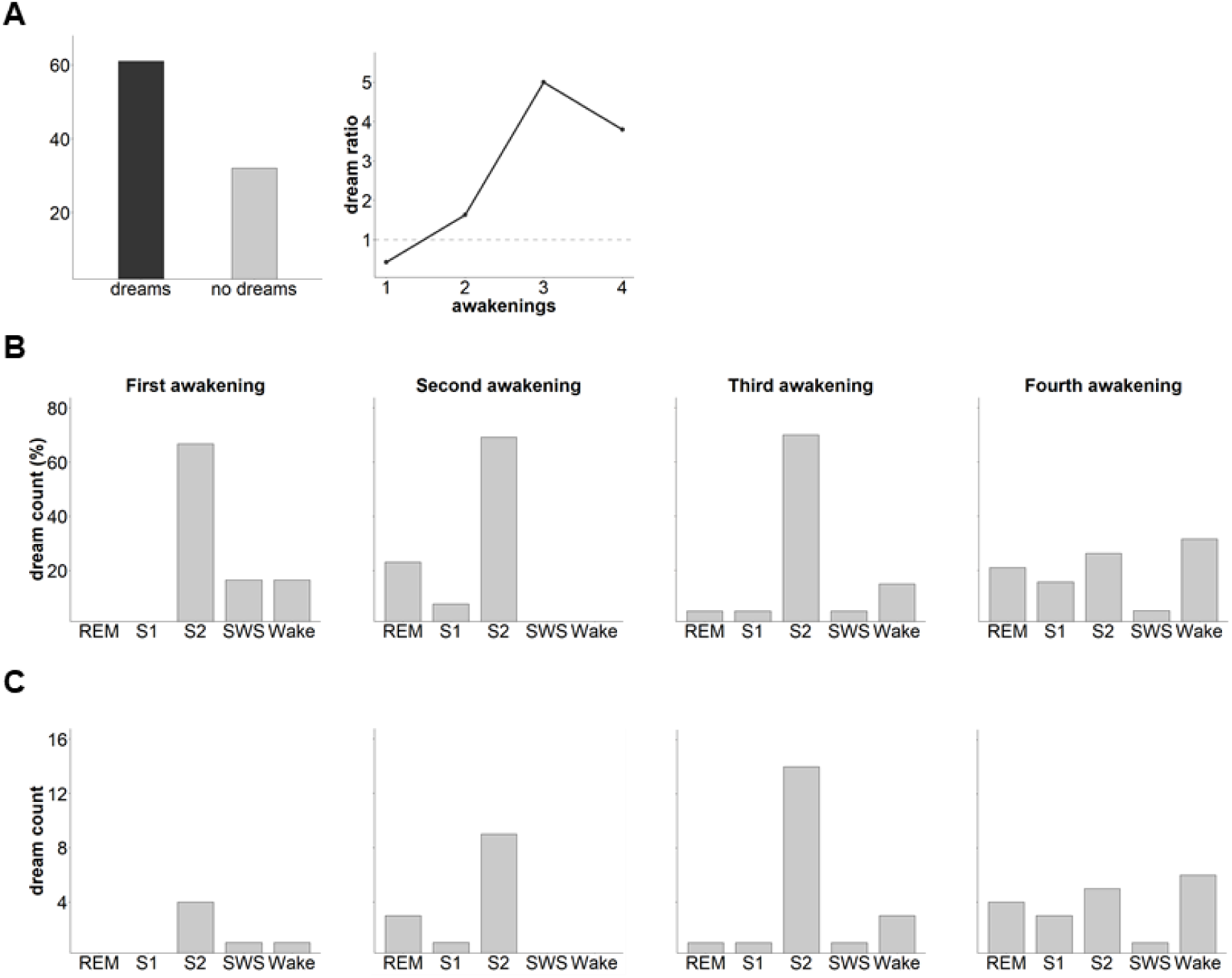
Distribution of collected dream reports and no dreams reported, across awakenings and sleep stages. **A.** Proportion of dream reports collected throughout the whole night in contrast with no dreams reported. The bar graph on the left displays the absolute numbers whereas the graph on the right depicts the ratio of dreams vs no dreams reported across awakenings. **B.** Last sleep stage experienced before reporting a dream, from the first to the fourth awakening, in percentage. **C.** Last sleep stage experienced before reporting a dream, from the first to the fourth awakening, absolute numbers.

**Figure S2.**
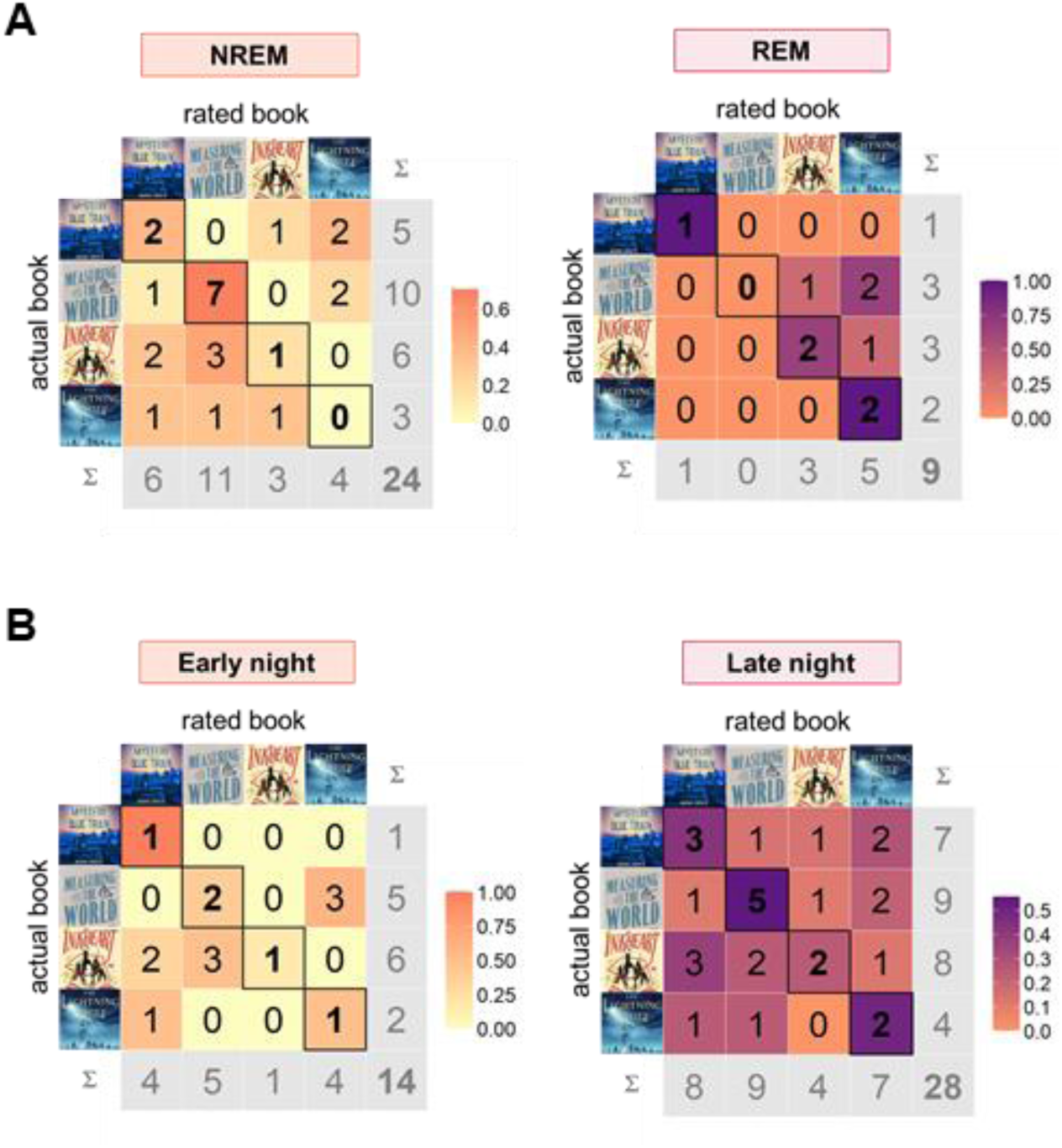
**Confusion matrix of listened vs. rated audiobook for combined ratings, night dynamics. A. NREM and REM sleep awakenings**. NREM: ĸ = 0.172, z = 1.43, p = 0.15; REM: ĸ = 0.41, z = 2.52, p = 0.012. Total number of cases: 24 and 9, 11 missing values in NREM and 2 in REM (ambiguous ratings). **B. Early and late awakenings**. Early awakenings: ĸ = 0.176, z = 1.4, p = 0.16; Late awakenings ĸ = 0.237, z = 2.22, p = 0.03. Total number of cases: 14 and 28, 5 missing values in early awakenings and 13 in late awakenings (ambiguous ratings, no concordance found between the three raters).

**Figure S3.**
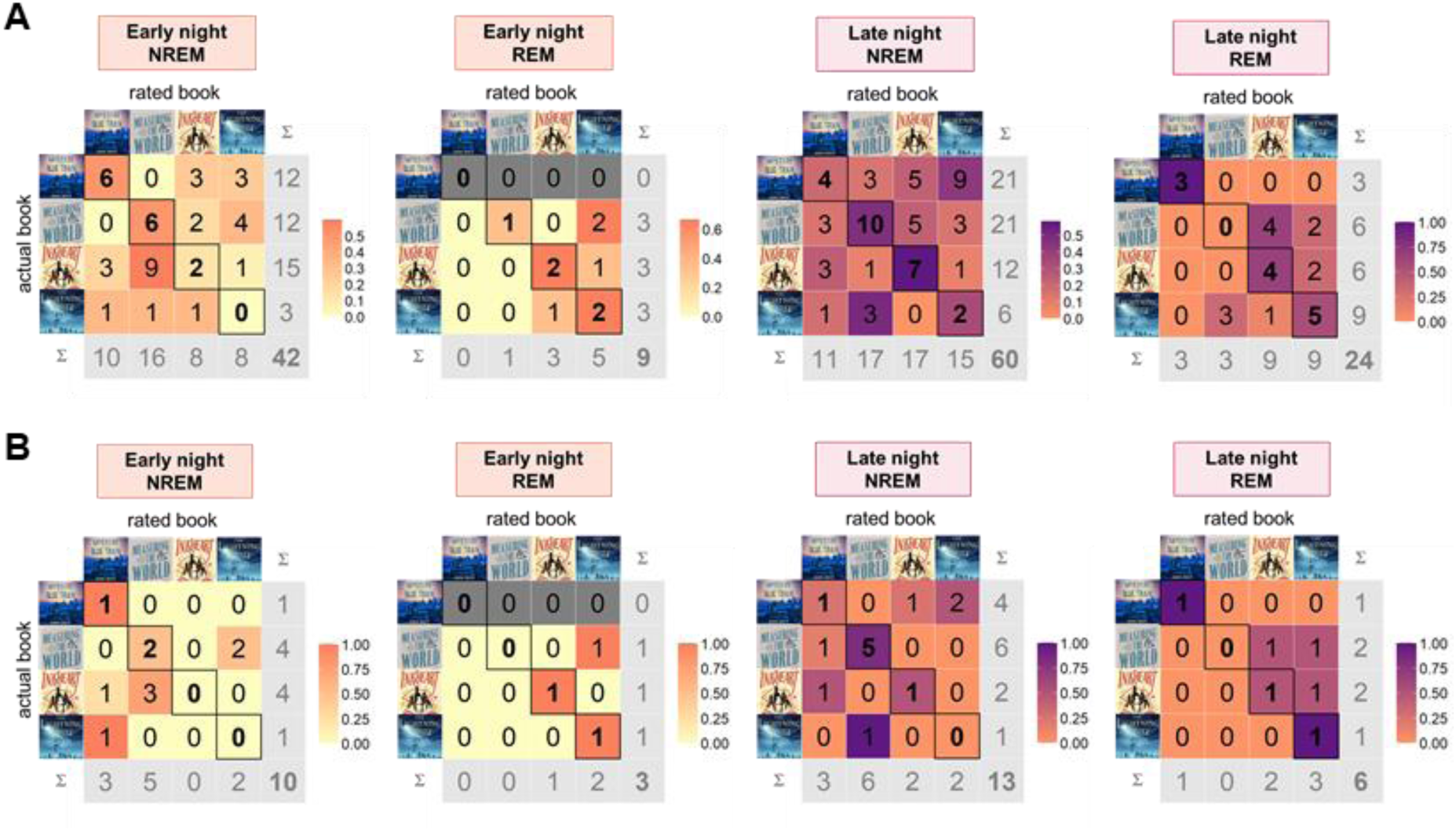
Confusion matrix of listened vs. rated audiobook for NREM and REM awakenings in individual and combined ratings, split by early and late night. **A**. **Individual rating.** From left to right: **Early night NREM,** ĸ = 0.101, z = 1.15, p = 0.25, total number of cases 42; **Early night REM**, ĸ = 0.333, z = 1.53, p = 0.13, total number of cases 9; **Late night NREM,** ĸ = 0.183, z = 2.58, p = 0.001, total number of cases 60; **Late night REM:** ĸ = 0.304, z = 2.52, p = 0.012, total number of cases 24. **B. Combined rating.** From left to right: **Early night NREM,** ĸ = 0.06, z = 0.467, p = 0.64, total number of cases 10, 3 missing values (ambiguous ratings, no concordance found between the three raters); **Early night REM**, ĸ = 0.5, z = 1.5, p = 0.13, total number of cases 3; **Late night NREM,** ĸ = 0.322, z = 1.88, p = 0.06, total number of cases 13, 8 missing values (ambiguous ratings, no concordance found between the three raters); **Late night REM:** ĸ = 0.357, z = 1.87, p = 0.06, total number of cases 6, 2 missing values (ambiguous ratings, no concordance found between the three raters);

**Figure S4.**
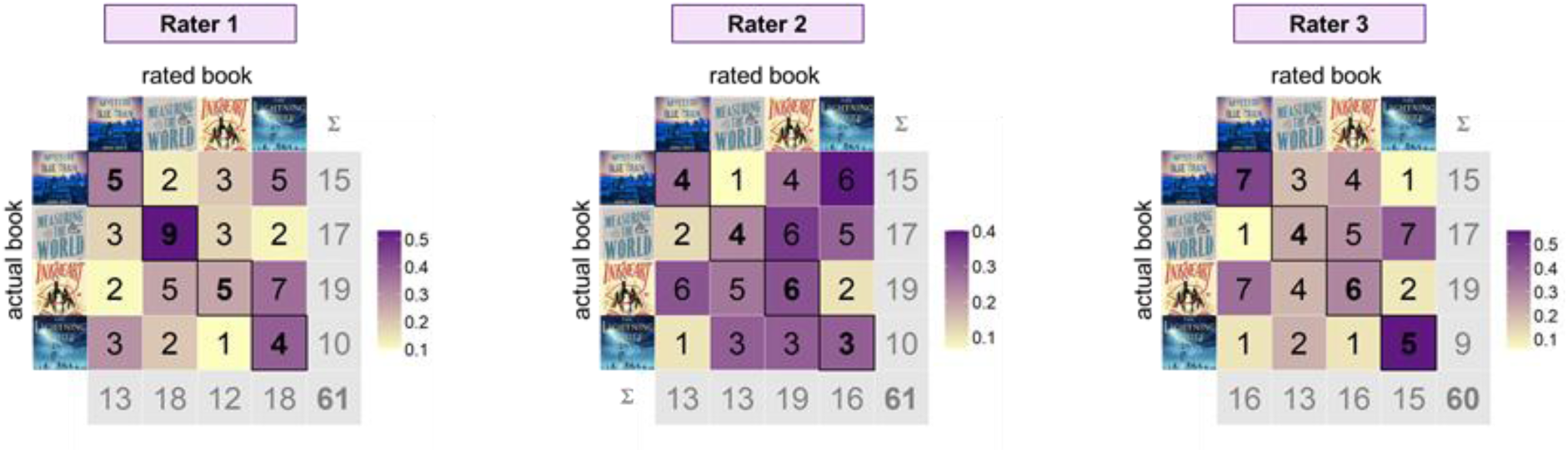
Dream rating analysis by single rater. Confusion matrix of listened against rated audiobook for each rater. **Rater 1:** ĸ = 0.176, z = 2.45, p = 0.014, total number of cases 61; **Rater 2:** ĸ = 0.036, z = 0.49, p = 0.62, total number of cases 61; **Rater 3:** ĸ = 0.156, z = 2.11, p=0.035, total number of cases 60, 1 missing value (missing rating).

**Figure S5.**
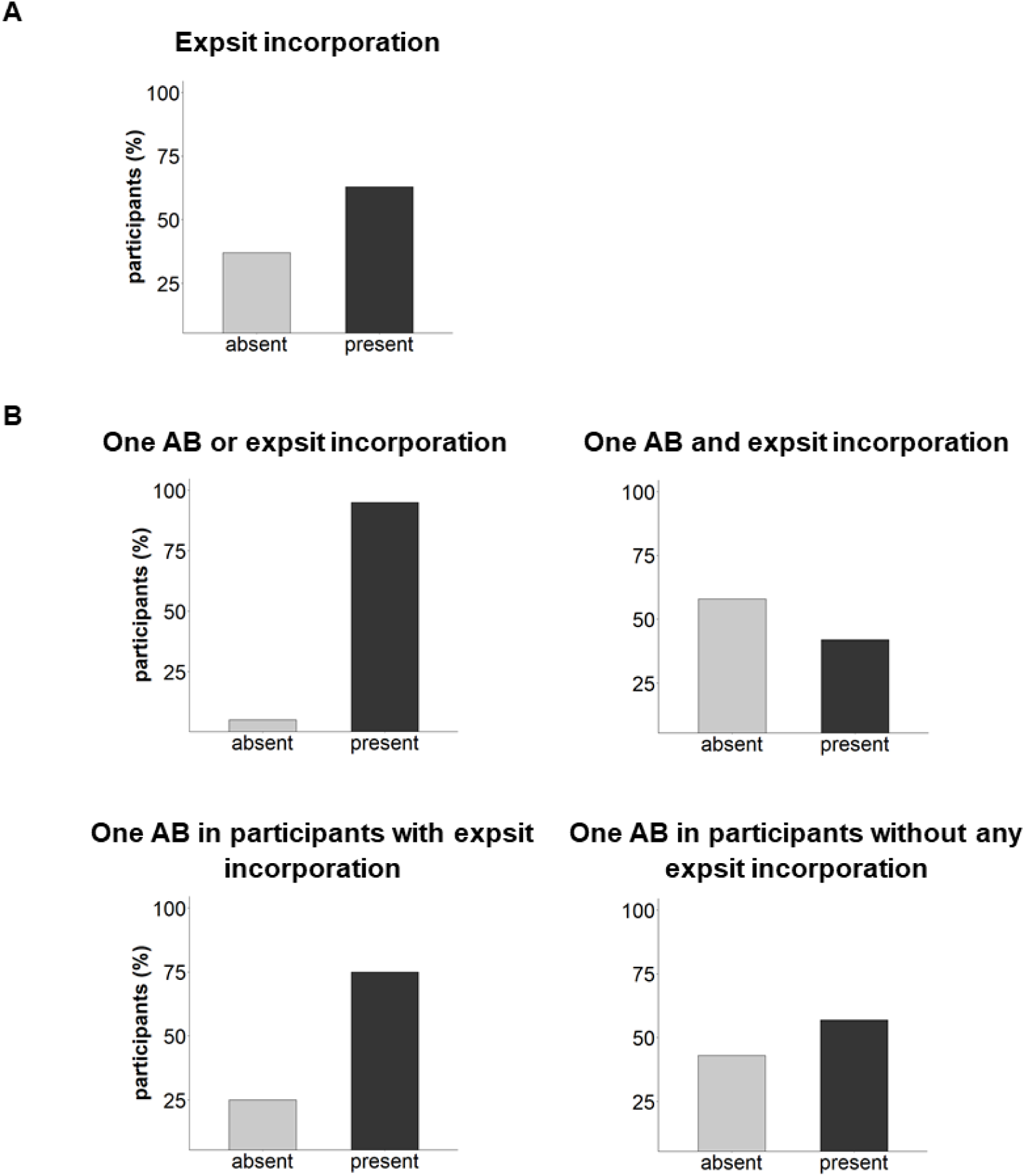
Distribution of incorporation of the experimental situation in dreams across participants. **A**. Proportion of participants with and without incorporation of the experimental situation (*expsit*) into dreams. **B. From top left to bottom right: (1)** Proportion of participants with at least one audiobook incorporation (*AB*) or experimental situation (*expsit*) incorporation into dreams; **(2)** Proportion of participants with at least one audiobook incorporation (*AB*) and experimental situation (*expsit*) incorporation into dreams; **(3)** Proportion of participants with and without audiobook incorporation (*AB*) from the pool of participants who reported at least one experimental situation (*expsit*) incorporation into dreams; **(4)** Proportion of participants with and without audiobook incorporation (*AB*) from the pool of participants who did not report any experimental situation (*expsit*) incorporation.

**Figure S6.**
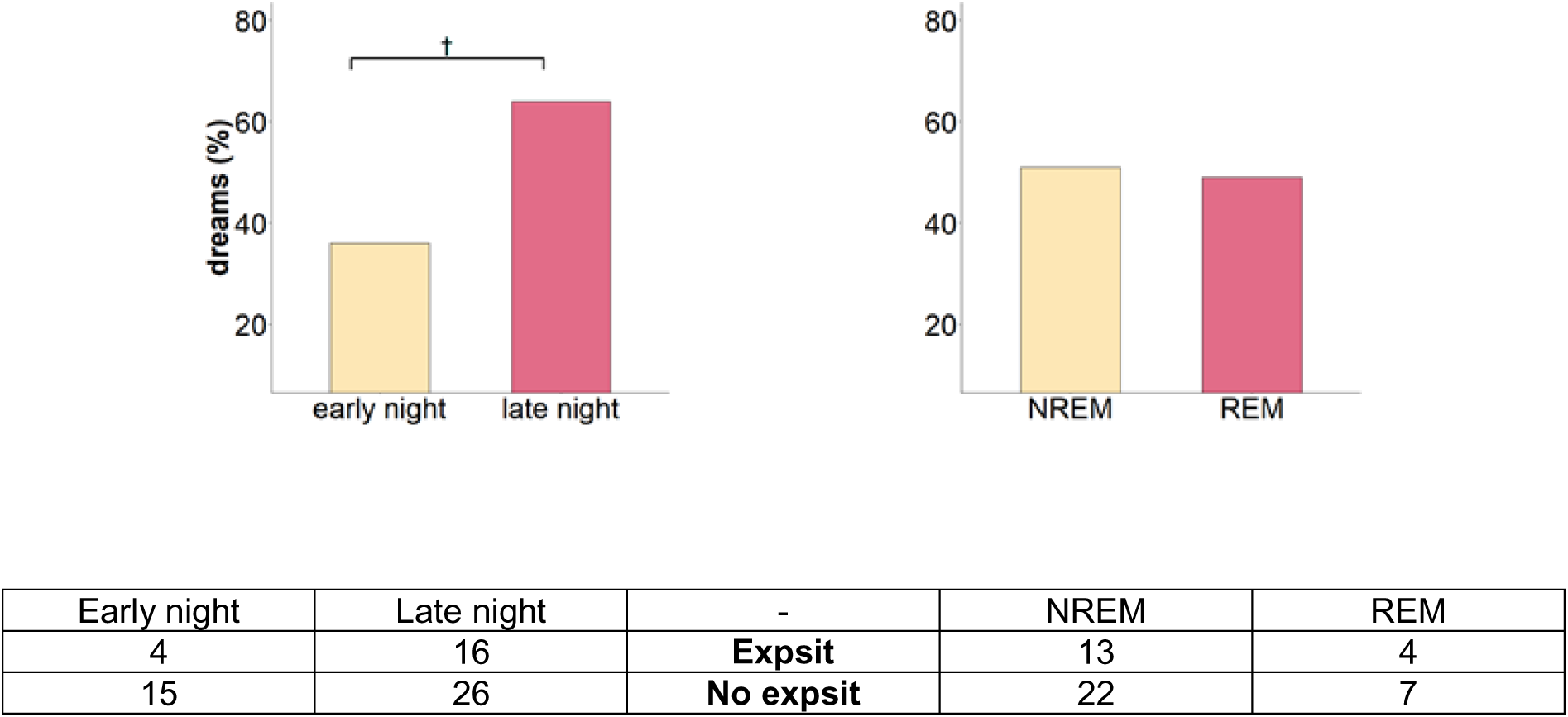
Distribution of dreams with and without incorporation of the experimental situation, night dynamics. Bar graph on the left: Distribution across early and late awakenings in percent. **Bar graph on the right:** Distribution across NREM and REM awakenings in percent. **Table:** Distribution across early and late awakenings (χ2(1, N_paired_ = 12) = 3.2, p =.07), as well as NREM and REM awakenings (χ2(1, N_paired_ = 7) = 0, p = 1) in absolute numbers. Cross indicates a trend in the group comparison (p < 0.1).

**Figure S7.**
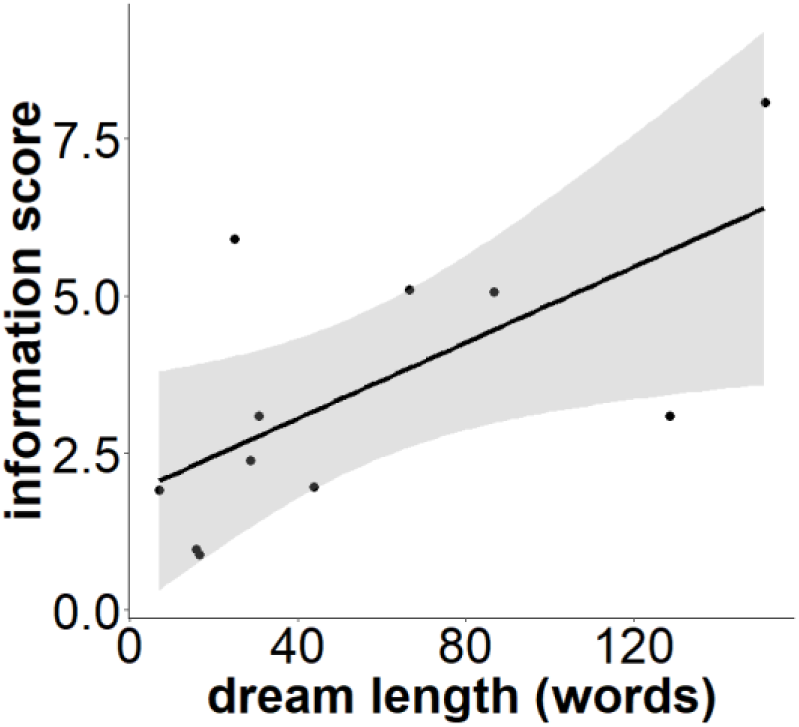
Relationship between dream length and audiobook incorporation. Relationship between the amount of words in dream reports and the amount of information reprocessed in dreams (Kendall’s Tau: τ = 0.51, z = 2.13, p = 0.03*, N = 11). * p < 0.05

**Figure S8.**
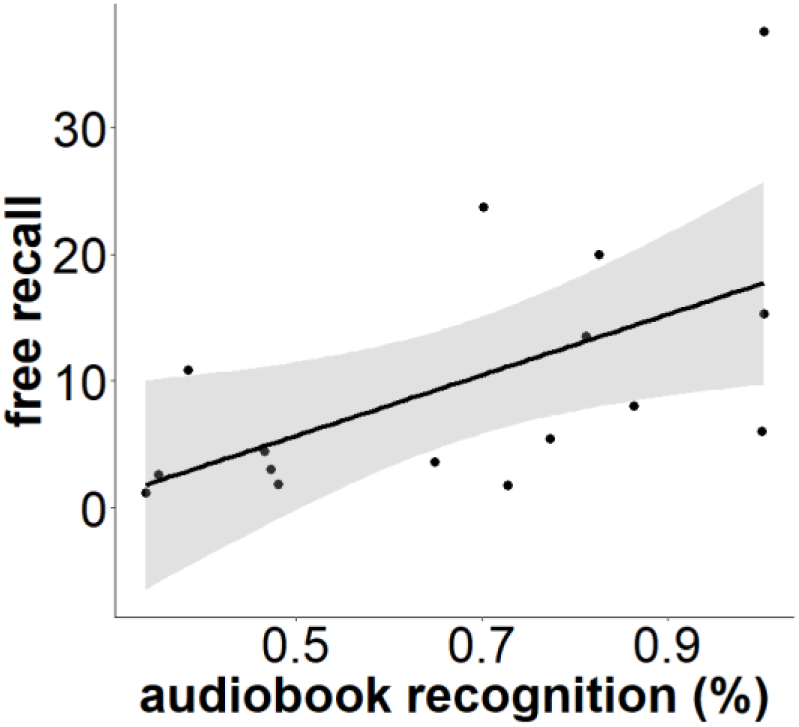
Relationship between audiobook recognition and free recall performance. Kendall’s Tau: τ = 0.483, z = 2.61, p = 0.0045** one-tailed, N = 16. ** p < 0.01

---

**Table S1.**
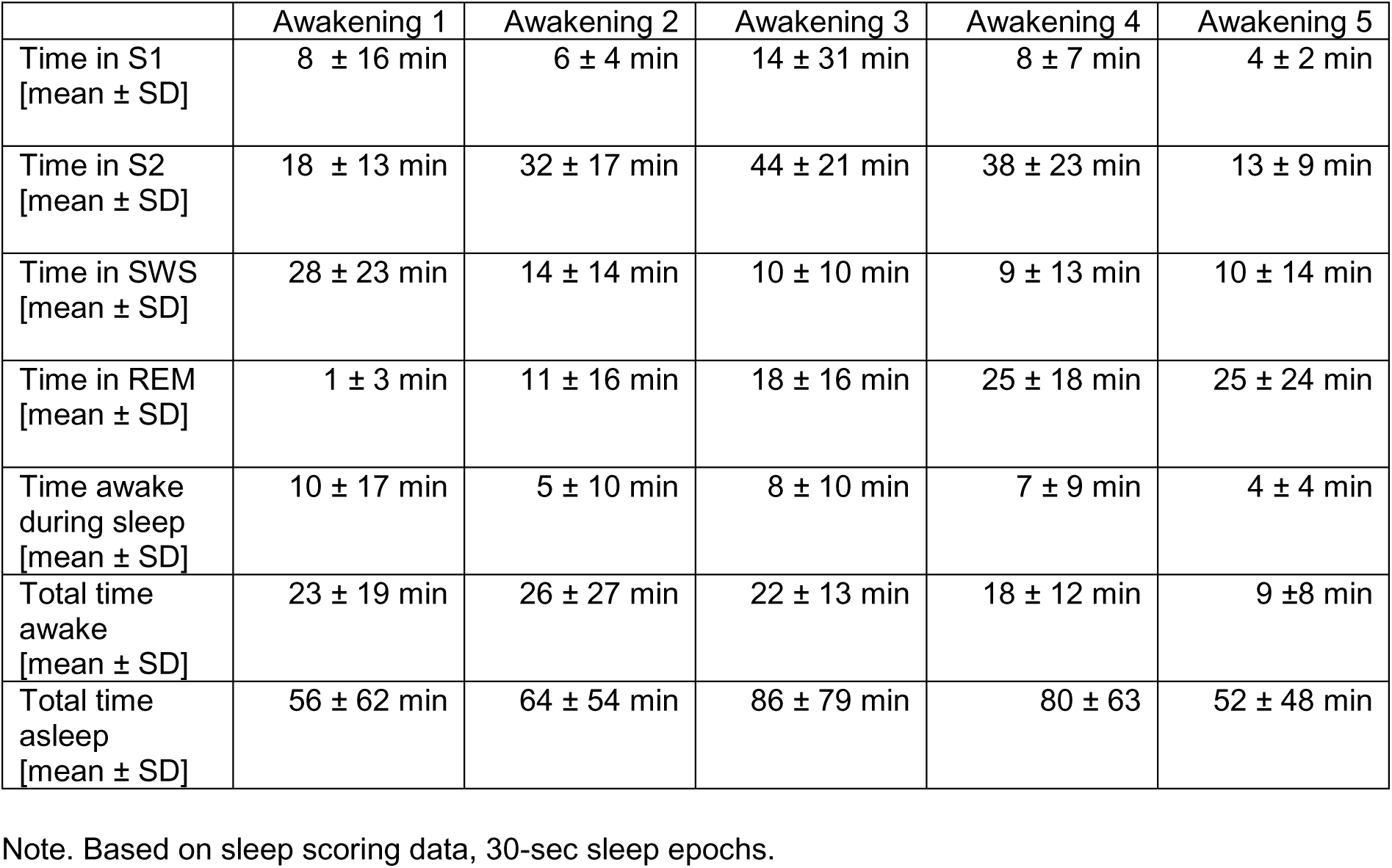
Time spent in sleep stage by awakening.

|  | Awakening 1 | Awakening 2 | Awakening 3 | Awakening 4 | Awakening 5 |
| --- | --- | --- | --- | --- | --- |
| Time in S1<br>[mean ± SD] | 8 ± 16 min | 6 ± 4 min | 14 ± 31 min | 8 ± 7 min | 4 ± 2 min |
| Time in S2<br>[mean ± SD] | 18 ± 13 min | 32 ± 17 min | 44 ± 21 min | 38 ± 23 min | 13 ± 9 min |
| Time in SWS<br>[mean ± SD] | 28 ± 23 min | 14 ± 14 min | 10 ± 10 min | 9 ± 13 min | 10 ± 14 min |
| Time in REM<br>[mean ± SD] | 1 ± 3 min | 11 ± 16 min | 18 ± 16 min | 25 ± 18 min | 25 ± 24 min |
| Time awake<br>during sleep<br>[mean ± SD] | 10 ± 17 min | 5 ± 10 min | 8 ± 10 min | 7 ± 9 min | 4 ± 4 min |
| Total time<br>awake<br>[mean ± SD] | 23 ± 19 min | 26 ± 27 min | 22 ± 13 min | 18 ± 12 min | 9 ± 8 min |
| Total time<br>asleep<br>[mean ± SD] | 56 ± 62 min | 64 ± 54 min | 86 ± 79 min | 80 ± 63 | 52 ± 48 min |

Note. Based on sleep scoring data, 30-sec sleep epochs.

**Table S2.**
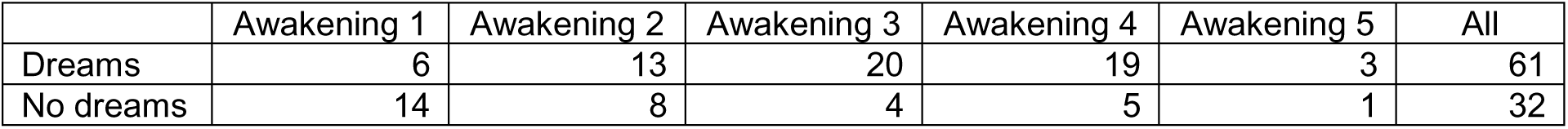
Number of dreams and no dreams by awakening.

|  | Awakening 1 | Awakening 2 | Awakening 3 | Awakening 4 | Awakening 5 | All |
| --- | --- | --- | --- | --- | --- | --- |
| Dreams | 6 | 13 | 20 | 19 | 3 | 61 |
| No dreams | 14 | 8 | 4 | 5 | 1 | 32 |

**Table S3.** Number of dreams and no dreams by participant.

| Participant | Dreams | No dreams |
| --- | --- | --- |
| 2 | 3 | 1 |
| 3 | 4 | 1 |
| 4 | 5 | 1 |
| 5 | 5 | 1 |
| 6 | 6 | 1 |
| 7 | 3 | 2 |
| 8 | 4 | 1 |
| 9 | 2 | 2 |
| 10 | 2 | 2 |
| 11 | 5 | 1 |
| 12 | 0 | 4 |
| 13 | 1 | 3 |
| 14 | 3 | 1 |
| 15 | 6 | 1 |
| 16 | 1 | 3 |
| 17 | 3 | 1 |
| 18 | 1 | 3 |
| 19 | 1 | 4 |
| 20 | 3 | 1 |
| 21 | 4 | 1 |
| average | 3.1 | 1.75 |
| SD(±) | 1.7 | 1.04 |

**Table S4.** Number of dreams and no dreams by last sleep stage prior to awakening.

|  | S1 | S2 | SWS | REM | WAKE |
| --- | --- | --- | --- | --- | --- |
| dreams | 5 | 32 | 3 | 11 | 10 |
| no dreams | 1 | 21 | 5 | 1 | 4 |
| total | 6 | 53 | 8 | 12 | 14 |

**Table S5.** Mean time spent in last sleep stage before reporting a dream.

|  | S1 | S2 | SWS | REM | WAKE |
| --- | --- | --- | --- | --- | --- |
| Avg time (min) | 6.40 | 8.72 | 12.17 | 24.55 | 15.60 |
| SD(±) | 7.8 | 8.69 | 12.97 | 18.15 | 11.25 |

**Table S6.** Distribution of last sleep stage by awakenings with dreams.

|  | Awakening 1 | Awakening 2 | Awakening 3 | Awakening 4 | Awakening 5 |
| --- | --- | --- | --- | --- | --- |
| S1 | 0 | 1 | 1 | 2 | 0 |
| S2 | 4 | 8 | 11 | 4 | 0 |
| SWS | 1 | 0 | 1 | 1 | 0 |
| REM | 0 | 3 | 1 | 3 | 2 |
| WAKE* | 1 | 0 | 2 | 4 | 0 |

\*In 6 instances out of 7, reports after wakefulness have been collected from late night awakenings, when participants already knew they would be asked for a dream upon multiple intervals, thus retaining their dream recall after spontaneously waking up before the planned awakening. These reports have been further evaluated to confirm that they resemble dream narratives and not waking thoughts or confabulations.

**Table S7a.** Indirect and direct audiobook incorporations in dream reports where all three raters agreed.

|  | Translated dream |
| --- | --- |
| Dream 1 | *"I tied ropes, I was alone on a sailing ship; there were also seals around; the sailing ship looked like a toy, an old, small Viking ship; there were also seals; I tied a rope; I was between ice, in cold water, there were seals on the ice; I tried to get away; I saw sails with ropes."* |
| Dream 2 | *"Someone said to me that an exchange to Brazil doesn't make sense, I should only go to Portugal; I dreamt that my experimenter showed it to me on the Iphone on a map, specifically."* |
| Dream 3 | *"I have the feeling that I dreamt about the audiobook, that the audiobook went on, but somehow it also transformed into TKKG; I always have this Elinor in my ear, that was the aunt's name, and that transformed into TKKG where people also had to flee in the night and hide from people, where threatening people were chasing them; also in a dark street; people, two people, one of whom was a boy and the other a girl, and I saw the whole thing from the perspective of the girl, but that wasn't me; I think they also had to run away from someone who wanted something from them and these people were very threatening. For example, the girl and the boy lay down on the floor to hide from them and then somehow an old woman came and bent down and meanly laughs: Hehehehe."* |
| Dream 4 | *"I was in an airplane with my father and he was still driving on the runway, then he said, I have to take off now or something and then he just drove faster but somehow he couldn't take off; then he let the plane roll out; I asked him: but, you can't actually fly and then he said, let's see: and it was at night and it was raining and there were people on the runway and they jumped away from him."* |
| Dream 5 | *"I had a huge fight with my boyfriend; we were running around the flat and yelling at each other; there was water damage in the flat; there were a bunch of strangers downstairs and somehow [the boyfriend] sent me away and I had to do it again; which is stupid because I never have to send strangers away from our flat; that really sucked."* |
| Dream 6 | *"What went through my mind was something about stolen diamonds and somehow that there was also a Chip and Chap episode along those lines."* |
| Dream 7 | *"I was in [city], at my parents'; briefly for background; my parents have a house and there is a flat next door that they have rented, there is a practice in there, you sometimes have to go over there to use the computer; that was the content of my dream that I have to go over there to the other buildings; I had to put on my shoes and get the key, that's what I did; the interesting thing was that I stand near the key board and think to myself, okay, I have to get the key and the thought made me pull it towards me; then I was downstairs and needed the shoes and I didn't just get them but I stood there with open hands and the thought was that the shoes should come to me and I already had them in my hands; I got dressed, then I went out, that is, I was in front of our house in [city]; there was a front garden and there was a dog, that half surprised me, because for one thing we used to have a dog, but I know that we don't have a dog at the moment; then this dog came running towards me, at first I had the impression that it was happy and positively minded; Nevertheless, he pulled me down to the ground, overpowered me so to speak, I noticed that from the perspective that I went down and the dog jumped on top of me; and that scared me so much that I actually thought I had woken up, I had this room around me and actually the feeling that I was screaming, but that nothing was coming out, so I couldn't hear anything; I then started to roll over, which probably isn't true, I don't think* |
|  | *so, but emotionally it was like this, I turned over here, lay on my stomach, then over here again in this direction, so relatively active, always trying to scream; and then something set in that I know from other dreams, namely that they are linked to a very real body perception; from that moment on it felt like the body perception now and I had the feeling that I was falling, it was nothing special but like in an elevator, this feeling of falling; it felt like I was falling through the floor here; I thought I was waking up in the dream and now I really woke up.”* |

**Table S7b.** Indirect and direct audiobook incorporations in dream reports where all three raters agreed: references to the audiobook.

|  | **Incorporation**<br>**(translated)** | **Title** | **Time of related**<br>**passage in the**<br>**audiobook** | **Listened**<br>**audiobook slot** |
| --- | --- | --- | --- | --- |
| Dream 1 | *"Humboldt is on [a] sailing ship."* | Measuring the World | 00:49:46 - 00:53:21 | 00:43:00 - 00:53:00 |
| Dream 2 | *"Map, travelling to faraway countries. South America, Portugal: Similar to Humboldt's travel stops."* | Measuring the World | 00:14:23;<br>00:46:45 - 00:56:36;<br>01:25:18; 01:26:46;<br>01:36:31 | 00:00:00 - 2:00:00 |
| Dream 3 | *"Elinor (explicit name mentioned), threat, running away, girl."* | Inkheart | 00:19:09;<br>00:20:09 - 00:34:01;<br>00:56:55 - 01:10:25;<br>01:16:09 - 01:23:02;<br>01:39:55 - 01:49:36;<br>01:55:10 - 02:00:43;<br>02:03:47 - 02:08:32;<br>02:12:29 - 02:19:00; | 1:17:00 - 2:19:00 |
| Dream 4 | *"The escape, the haste, the police and the relationship with the father. "* | Inkheart | General theme of book (night, haste, flight, father) | 00:29:00 - 1:02:00 |
| Dream 5 | *"Water, fountain; Percy causes water damage (own apartment mixed with elements from Percy). "* | Percy Jackson | General theme of the book (water) | 1:06:00 - 1:25:00 |
| Dream 6 | *"Jewels, diamonds; diamond theft (or at the time indicated, the attempted theft); The theme song of Chip 'n' Dale has the lyrics 'Robbers, thieves are on a crooks' tour.'"* | The Mystery of the Blue Train | 00:00:00 - 00:12:39<br>00:29:21 | 00:25:00 – 00:56:00 |
| Dream 7 | *"Magic: movement of objects through the power of thought, attacked by a dog."* | Percy Jackson | General theme of the book<br>1:51:39 (anticipation of dog attack) | 1:33:00 - 1:55:00 |

**Table S7c.** Indirect and direct audiobook incorporations in dream reports where all three raters agreed: references to the audiobook.

|  | Audiobook passage<br>(translated) |
| --- | --- |
| Dream 1<br>(Measuring<br>the World) | **"They took the first frigate** that departed from La Coruña for the tropics. The wind blew sharply from the west, the sea was rough. **Humboldt sat in a folding chair on deck.** He felt freer than ever before. Luckily, he wrote in his journal, he never got seasick. (...) Shortly before reaching Tenerife, they sighted a sea monster. In the distance, almost transparent against the horizon, a serpentine body rose from the water, formed two ring-like coils, and looked over at them with gemstone eyes clearly visible through the telescope. Fine, beard-like threads hung from its mouth. Just seconds after it submerged again, everyone believed they'd imagined it." |
| Dream 2<br>(Measuring<br>the World) | "Alexander von Humboldt was famous throughout Europe for an **expedition to the tropics** that he had undertaken twenty-five years earlier. He had been in **New Spain, New Granada, New Barcelona, New Andalusia,** and the United States, had discovered the natural canal between the **Orinoco and the Amazon.** (...) And now what? Where else, replied Humboldt, but to Spain! (...) From then on, they traveled at night so he could take measurements undisturbed. **The plane coordinates had to be determined more precisely** than had been done so far. **The maps of Spain** were not accurate. One wanted to know where one was riding. But we do know, cried Bonpland. Here's the highway, and it leads to Madrid. What more do you need! It's not about the road, replied Humboldt. It's about the principle. (...) According to old reports, there was a canal between the Orinoco and Amazon rivers. He would find the canal and solve the mystery. Ah, said Bonpland. A canal. **Humboldt didn't like this attitude. Always complaints, always objections.** Was a bit of enthusiasm too much to ask?" |
| Dream 3<br>(Inkheart) | **"Elinor's hair was already going gray,** and though she had pinned it up, strands were hanging loose everywhere, as if she'd done it hastily and with impatience. **Elinor didn't look like someone who spent much time in front of the mirror.** »Good heavens, Mortimer! Well, if that isn't a surprise!« she said, skipping any formal greeting. »Where did you come from?« Her voice sounded gruff, but her face couldn't completely hide that she was pleased to see Mo. »Hello, Elinor,« said Mo, placing a hand on Meggie's shoulder. »Do you remember Meggie? She's grown quite a bit, as you can see.« **Elinor gave Meggie a quick, puzzled glance.** (...) [Meggie] She thought she heard voices, louder than Dustfinger's music, men's voices, and **a terrible fear began to spread within her,** as black and foreign as the night when Dustfinger had stood outside in the yard. When she jumped up, Dustfinger dropped a burning torch, and it fell to the grass. Quickly, he stomped out the fire before it could spread, then followed Meggie's gaze and saw her looking toward the house, without saying a word. **But Meggie ran.** (...) **She kept running, limping and sobbing, until she stood in front of the large iron gate. But the street behind it was empty. And Mo was gone.** (...) Elinor stood in the brightly lit front door as Meggie returned. She had thrown a coat over her nightdress. The night was warm, but now a cold wind was blowing up from the lake. **How desperate the girl looked –** so lost. Elinor remembered the feeling. There was no worse feeling. 'They've taken him!' Meggie's voice nearly choked on her helpless rage. (...) When she woke up, suddenly, with her heart pounding and her hair damp with sweat, **everything was immediately back: the men, Mo's voice, and the empty street.** I'll go look for him, Meggie thought. Yes, I will." |
| Dream 4<br>(Inkheart) | "And there she saw him. **The darkness was pale from the rain,** and the stranger [Dustfinger] was barely more than a shadow. Only his face glowed toward Meggie. His hair stuck to his wet forehead. The rain was dripping down on him, but he paid it no mind. He stood there motionless, his arms crossed over his chest, as if that were the only way to keep himself even a little warm. He stared across at their house. I have to wake Mo!, Meggie thought. (...) »But he'll get it one way or another! I'm telling you again—they're on your trail.« »That wouldn't be the first time. So far, I've always managed to shake them off.« »Oh really? And how long do you think that'll keep working? And what about your daughter? Are you going to tell me she enjoys moving from place to place all the time? Believe me, I know what I'm talking about.« (...) It was just getting light when Meggie was jolted out of sleep. Over the fields, the night faded, as if the rain had washed away the hem of its dress. On the alarm clock, it was just before five, and Meggie wanted to turn over and sleep on, but suddenly she felt that someone was in the room. Startled, she sat up and saw Mo standing in front of her open wardrobe. »Good morning!« he said, as he put her favorite sweater into a suitcase. »Sorry, I know it's very early, but we have to travel. How about hot chocolate for breakfast?« Meggie nodded sleepily. Outside, the birds were chirping as if they had been awake for hours. Mo threw two more pairs of her pants into the suitcase, closed it, and carried it to the door. »Put something warm on,« he said. »It's cold outside.« »Where are we going?« Meggie asked. (...) **Yes. Meggie loved the bus, but on that morning, she hesitated to climb inside. When Mo went back to the house to lock the door, she suddenly had the feeling that they would never return, that this trip would be different from all the others, that they would keep** |
|  | *driving and driving, fleeing from something that had no name. At least none that Mo had told her. »So, off to the south!« he said, as he squeezed behind the steering wheel. And so they set off – without saying goodbye to anyone, on a much too early morning that smelled of rain. But at the gate, Dustfinger was already waiting for them.”* |
| Dream 5<br>(Percy Jackson) | *“I was just about to unpack my sandwich when Nancy Bobofit showed up in front of me with her ugly friends and dumped the remains of her lunch into Grover’s lap. »Yikes.« She grinned at me, showing her crooked teeth. I tried to stay completely cool. The school psychologist had advised me that a million times. »Count to ten, then you’ll get your anger under control.” But I was so angry that I couldn’t think anymore. A wave seemed to roar in my ears. I can’t remember touching her, but suddenly Nancy was sitting in the fountain and screaming: »Percy pushed me!” Immediately Mrs. Dodds was standing next to us. Some of the others were whispering: »Did you see ...« »... the water ...« »... as if it had grabbed her ...« I had no idea what they were talking about. I only knew that I was in trouble again.”* |
| Dream 6<br>(Mystery of the Blue Train) | *“The Russian looked disturbed and worried. »Are you sure – that the package is safe? No one has tampered with it? There’s been so much talk – far too much talk.« He started chewing on his nails again. »Check for yourself.« She leaned over to the fireplace and skilfully pushed the coals aside. Beneath them lay crumpled paper bundles; from among them, she took a long package, wrapped in greasy newspaper, and handed it to the man. (...)»I must apologize for – for this unusual meeting place. But confidentiality is essential. I – I can’t afford to be linked to this business. » »I understand,” said the American politely. »I have your word, don’t I, that no detail of this transaction will be made public? That’s one of the conditions of sale.” The American nodded. »We agreed on that,« he said evenly. »Perhaps you could show me the goods now. » »You have the money with you – in cash?“ »Yes,« the other man replied. (...) After a brief hesitation, Krassnine gestured toward the package on the table. **The American took it and unwrapped the packing paper. He moved to a small lamp and inspected the contents thoroughly. He seemed satisfied, pulled a thick leather wallet from his pocket, and took out a bundle of banknotes.** He handed them to the Russian, who counted them carefully. (...) **The secretary gasped for air. On the dirty white pad, the stones glowed like blood.** »My God! Sir,« said Knighton. »Are they – are they real? » Van Aldin let out a quiet, amused chuckle. »Doesn’t surprise me that you ask. Among these rubies are the three largest in the world. Catherine of Russia wore them, Knighton. **The one in the middle is known as the Heart of Fire. It is perfect – not the slightest flaw.** » »But, they must be worth a fortune. »”* |
| Dream 7<br>(Percy Jackson) | *“That evening, things were much more exciting than usual after dinner. It was Flag Capture Day. (...) »Heroes and heroines!« he called out. »You know the rules. The stream is the boundary. The entire forest is allowed. **All magical things are permitted.** (...) »So, what do we do now?“ I asked. »Can you lend me any **magical tools?**« (...) Then, very close by, **I heard a faint growl. I instinctively raised my shield; I had the feeling that something was sneaking up on me. Then the growling stopped. I felt that the something was retreating.**”* |

Only the most relevant audiobook passages are included in the table. Gray-shaded sections refer to earlier segments and are provided for additional context. The most pertinent parts of the text are highlighted in bold.

**Table S8a.** Influence of early vs late night on memory benefit, linear regression.

| Predictor | $\beta$<br>(Estimate) | t(10) | p-value | 95% CI (Lower) | 95% CI (Upper) |
| --- | --- | --- | --- | --- | --- |
| Intercept<br>(early night) | 0.60 | 1.89 | 0.09 | -0.11 | 1.32 |
| Info score | 0.07 | 0.79 | 0.45 | -0.12 | 0.25 |
| Night segment (late<br>night) | -0.34 | -0.94 | 0.37 | -1.14 | 0.46 |
| Info score x Night<br>segment (late night) | 0.006 | 0.07 | 0.95 | -0.2 | 0.21 |

Summary:

**Table S8b.** Influence of last sleep stage prior to awakening on memory benefit, linear regression.

| Predictor | $\beta$<br>(Estimate) | t(8) | p-value | 95% CI (Lower) | 95% CI (Upper) |
| --- | --- | --- | --- | --- | --- |
| Intercept<br>(last sleep stage<br>NREM) | 0.08 | 0.53 | 0.61 | -0.26 | 0.42 |
| Info score | 0.11 | 2.57 | 0.03 | 0.01 | 0.21 |
| Last sleep stage (REM) | 0.63 | 2.48 | 0.04 | 0.04 | 1.21 |
| Info score x Last sleep<br>stage segment (REM) | -0.09 | -1.57 | 0.15 | -0.23 | 0.04 |

Summary:
F(3, 8) = 5.02, p = 0.030, R<sup>2</sup> = 0.65, Adjusted R<sup>2</sup> = 0.52

**Table S9.** Incorporation of the experimental situation.

|  | Translated dream |
| --- | --- |
| Absent | *„I dreamt that I was playing the violin; and all the time there was a song of chanson called Poeme playing.“* |
| Present - implicit | *“And then I was connected to things, which was no surprise; the whole world was connected to things with cables and each had a number; I think what you said yesterday surfaced; that it always has a number and you have to pull on it and each number belongs in a clip and in my dream, everyone in the world had such a thing and they were all connected to a mother cable thing and all lived underground in the dark. It was kind of a dark dream, like a dome under the earth and there were an exorbitant number of people, an insane number of people, and they all had a stick thing on their heads, from which a tube went quite far upwards; there was a huge tube from the dome somewhere, I don't know. We definitely had white full-body suits on.”* |
| Present- explicit | *“We were here and a sleep study was carried out, one person was definitely there, of whom I know who he is, he was part of our own study; he then carried out tests but everything went wrong, he kept coming and asking something; in the room where the computers are, there were several more people, I think you [experimenter 1] were there too and [experimenter 2] and someone I don't know; the last thing was that this person from my study came in and sprayed with tear gas, he came in and there was a lot of fog and he asked where something was; he had asked several questions, he came two or three times.”* |

## REFERENCES

1. Crick F & Mitchinson G. The function of dream sleep. Nature. 1983; Jul 14-20;304(5922): 111-4. 10.1038/304111a0.

2. Revonsuo A. The reinterpretation of dreams: An evolutionary hypothesis of the function of dreaming. Behav Brain Sci. 2000; Dec 23(6): 877-901. doi:10.1017/S0140525X00004015.

3. Freud S. The Interpretation of Dreams. 1900.

4. Hall C & Nordby V. The individual and his dreams. New American Library, New York. 1972.

5. Domhoff G W. Finding Meaning in Dreams: A Quantitative Approach. Plenum Press. 1996. 10.1007/978-1-4899-0298-6.

6. Strauch I & Meyer B. In Search of Dreams. Results of Experimental Dream Research. State University of New York Press. 1996.

7. Schredl M. Dream Recall: Research, Clinical Implications and Future Directions. Sleep and Hypnosis. 1999; 1(2): 72–81.

8. Schredl M. Factors Affecting the Continuity Between Waking and Dreaming: Emotional Intensity and Emotional Tone of the Waking-Life Event. Sleep and Hypnosis. 2006; 8(1): 1–5.

9. Malinowski J & Horton C L. Evidence for the Preferential Incorporation of Emotional Waking-Life Experiences Into Dreams. Dreaming. 2014; 24(1): 18–31. 10.1037/a0036017.

10. Breger L, Hunter I & Lane R W. The effect of stress on dreams. Psychological issues. 1971;7(3): 213.

11. Gorgoni M, Scarpelli S, Alfonsi V, Annarumma L, Cordone S, Stravolo S, De Gennaro L. Pandemic dreams: quantitative and qualitative features of the oneiric activity during the lockdown due to COVID-19 in Italy. Sleep Medicine. 2021; 81: 20–32. doi: 10.1016/j.sleep.2021.02.006.

12. Stickgold R, Hobson J A, Fosse R, Fosse M. Sleep, Learning, and Dreams: Off-line Memory Reprocessing. Science. 2001; 294: 1052–1057. 10.1126/science.1063530.

13. Wamsley E J & Stickgold R. Memory, Sleep, and Dreaming: Experiencing Consolidation. Sleep Medicine Clinics. 2011; 6(1): 97–108. 10.1016/j.jsmc.2010.12.008.

14. Wamsley E J. Dreaming and Offline Memory Consolidation. Curr Neurol Neurosci Rep. 2014; 14 (3): 433. doi: 10.1007/s11910-013-0433-5.

15. Pavlides C & Winson J. Influences of hippocampal place cell firing in the awake state on the activity of these cells during subsequent sleep episodes. J. Neurosci. 1989; 9 (8): 2907–2918. doi: 10.1523/JNEUROSCI.09-08-02907.1989.

16. Wilson M A & McNaughton B L. Reactivation of hippocampal ensemble memories during sleep. Science. 1994; 265: 676–679. 10.1126/science.8036517.

17. Ji D & Wilson M A. Coordinated memory replay in the visual cortex and hippocampus during sleep. Nat Neurosci. 2007; 10 (1): 100–107. doi: 10.1038/nn1825.

18. Maquet P, Laureys S, Peigneux P, et al. Experience-dependent changes in cerebral activation during human REM sleep. Nature neuroscience. 2000; 3 (8): 831–6. doi: 10.1038/77744.

19. Peigneux P, Laureys S, Fuchs S, et al. Are Spatial Memories Strengthened in the Human Hippocampus during Slow Wave Sleep? Neuron. 2004; 28, 44 (3): 535-45. doi:10.1016/j.neuron.2004.10.007.

20. Deuker L, Olligs J, Fell J, et al. Memory Consolidation by Replay of Stimulus-Specific Neural Activity. J Neurosci. 2013; Dec 4, 33 (49): 19373-83. doi: 10.1523/JNEUROSCI.0414-13.2013.

21. Schönauer M, Alizadeh S, Jamalabadi H, Abraham A, Pawlizki A, Gais S. Decoding material specific memory reprocessing during sleep in humans. Nat Commun. 2017; May 17, 8: 15404. doi: 10.1038/ncomms15404.

22. Schreiner T, Petzka M, Staudigl T, Staresina B P. Endogenous memory reactivation during sleep in humans is clocked by slow oscillation-spindle complexes. Nat Commun. 2021; 12: 3112. doi: 10.1038/s41467-021-23520-2.

23. Stickgold R, Malia A, Maguire D, Roddenberry D, O’Connor M. Replaying the game: hypnagogic images in normals and amnesics. Science. 2000; 290: 350–353. 10.1126/science.290.5490.350.

24. Kusse C, Shaffii-LE Bourdiec A, Schrouff J, Matarazzo L & Maquet P. Experience-dependent induction of hypnagogic images during daytime naps: a combined behavioural and EEG study. J Sleep Res. 2012; 21 (1): 10–20. doi: 10.1111/j.1365-2869.2011.00939.x.

25. Wamsley E J, Perry K, Djonlagic I, Reaven L B & Stickgold R. Cognitive replay of visuomotor learning at sleep onset: temporal dynamics and relationship to task performance. Sleep. 2010; 33 (1): 59–68. doi: 10.1093/sleep/33.1.59.

26. Wamsley E J, Tucker M, Payne J D, Benavides J A & Stickgold R. Dreaming of a learning task is associated with enhanced sleep-dependent memory consolidation. Curr. Biol. 2010; 20 (9): 850–855. doi: 10.1016/j.cub.2010.03.027.

27. Schoch S F, Cordi M J, Schredl M & Rasch B. The effect of dream report collection and dream incorporation on memory consolidation during sleep. J Sleep Res. 2019; 28 (1). doi:10.1111/jsr.12754.

28. Plailly J, Villalba M, Vallat R, Nicolas A, Ruby P. Incorporation of fragmented visuo-olfactory episodic memory into dreams and its association with memory performance. Sci Rep. 2019, 30, 9 (1): 15687. doi: 10.1038/s41598-019-51497-y.

29. Wamsley E J & Stickgold R. Dreaming of a learning task is associated with enhanced memory consolidation: Replication in an overnight sleep study. J Sleep Res. 2019; 28. doi: 10.1111/jsr.12749.

30. De Koninck J, Prévost F, Lortie-Lussier M. Vertical inversion of the visual field and REM sleep mentation. J. Sleep Res. 1996; 5 (1): 16–20. doi: 10.1046/j.1365-2869.1996.00001.x.

31. Cipolli C, Fagioli I, Mazzetti M, Tuozzi G. Incorporation of presleep stimuli into dream contents: evidence for a consolidation effect on declarative knowledge during REM sleep? J. Sleep Res. 2004; 13 (4): 317–26. doi: 10.1111/j.1365-2869.2004.00420.x.

32. Schredl M & Erlacher D. Is sleep-dependent memory consolidation of a visuo-motor task related to dream content? International Journal of Dream Research. 2010; 3 (1): 74–79.

33. Stamm A W, Nguyen N D, Seicol B J, et al. *EJ.* Negative reinforcement impairs overnight memory consolidation. Learn Mem. 2014; 15, 21 (11). doi: 10.1101/lm.035196.114.

34. Nefjodov I, Winkler A & Erlacher D. Balancing in dreams: Effects of playing games on the Wii balance board on dream content. International Journal of Dream Research. 2016; 9 (1): 89–92. 10.11588/ijodr.2016.1.29034.

35. Wamsley E J, Hamilton K, Graveline Y, Manceor S, Parr E. Test Expectation Enhances Memory Consolidation across Both Sleep and Wake. PLoS One. 2016; 19, 11 (10): e0165141. doi: 10.1371/journal.pone.0165141.

36. Fogel S M, Ray L B, Sergeeva V, De Koninck J, Owen A M. A Novel Approach to Dream Content Analysis Reveals Links Between Learning-Related Dream Incorporation and Cognitive Abilities. Frontiers in Psychology. 2018; 6, 9: 1398. doi: 10.3389/fpsyg.2018.01398.

37. Carr M F, Jadhav S P, Frank L M. Hippocampal replay in the awake state: a potential substrate for memory consolidation and retrieval. Nature Neurosci. 2011; 14 (2): 147–53. doi: 10.1038/nn.2732.

38. Nádasdy Z, Hirase H, Czurkó A, Csicsvari J & Buzsáki G. Replay and Time Compression of Recurring Spike Sequences in the Hippocampus. The Journal of Neuroscience. 1999; 19: 9497 9507. doi: 10.1523/JNEUROSCI.19-21-09497.1999.

39. Buzsaki G. The Hippocampo-Neocortical Dialogue. Cereb Cortex. 1996; 6 (2): 81–92. 10.1093/cercor/6.2.81.

40. Baylor G W & Cavallero C. Memory Sources Associated with REM and NREM Dream Reports Throughout the Night: A New Look at the Data. Sleep. 2001; 15, 24 (2): 165-70.

41. Fosse M J, Fosse R, Hobson J A & Stickgold R J. Dreaming and Episodic Memory: A Functional Dissociation? Journal of Cognitive Neuroscience. 2003; 15 (1): 1–9. doi:10.1162/089892903321107774.

42. Malinowski J E. & Horton C L. Memory sources of dreams: the incorporation of autobiographical rather than episodic experiences. J Sleep Res. 2014; 23 (4): 441–447. doi: 10.1111/jsr.12134.

43. Torda C. Dreams of subjects with bilateral hippocampal lesions. Acta Psychiatr Scand. 1969; 45 (3): 277–88. doi: 10.1111/j.1600-0447.1969.tb07128.x.

44. Nielsen T A & Stenstrom P. What are the memory sources of dreaming? Nature. 2005; 27, 437 (7063): 1286-9. doi: 10.1038/nature04288.

45. Spano G, Pizzamiglio G, McCormick C, et al. Dreaming with hippocampal damage. Elife. 2020; 8,9: e56211. doi: 10.7554/eLife.56211.

46. Schneider L. Anatomy and Physiology of Normal Sleep. Sleep and Neurologic Disease. 2017; (1): 1–28. 10.1016/B978-0-12-804074-4.00001-7.

47. Aserinsky E & Kleitman N. Regularly Occurring Periods of Eye Motility, and Concomitant Phenomena, During Sleep. Science. 1953; 4, 118 (3062): 273-4. doi:10.1126/science.118.3062.273.

48. Dement W. Cyclic variations in EEG during sleep and their relation to eye movements, body motility, and dreaming. Electroencephalogr Clin Neurophysiol. 1957; 9 (4): 673–90. doi:10.1016/0013-4694(57)90088-3.

49. Foulkes D. Dream reports from different stages of sleep. J Abnorm Soc Psychol. 1962; 65: 14–25. doi: 10.1037/h0040431.

50. Brown JN, Cartwright RD. Locating NREM dreaming through instrumental responses. Psychophysiology. 1978; 15 (1): 35–9. doi: 10.1111/j.1469-8986.1978.tb01330.x.

51. Rechtschaffen A, Verdone P & Wheaton J V. Reports of Mental Activity during Sleep. Canadian Psychiatric Association Journal. 1963; 8 (6): 409–414. doi: 10.1177/070674376300800612.

52. Monroe L J, Rechtschaffen A, Foulkes W D & Jensen J. Discriminability of REM and NREM reports. Journal of Personality and Social Psychology. 1965; 2 (3): 456–460. https://psycnet.apa.org/doi/10.1037/h0022218.

53. Cavallero C, Cicogna P, Natale V, Occhionero M & Zito A. Slow wave sleep dreaming. Sleep. 1992; 15 (6): 562–566. doi: 10.1093/sleep/15.6.562.

54. Antrobus J, Kondo T, Reinsel R, Fein G. Dreaming the late morning: summation of REM and diurnal cortical activation. Consciousness and Cognition. 1995; 4 (3): 275–299.10.1006/ccog.1995.1039.

55. Pivik T & Foulkes W D. NREM mentation: Relation to personality, orientation time, and time of night. Journal of Consulting and Clinical Psychology. 1968; 32 (2): 144–151. https://psycnet.apa.org/doi/10.1037/h0025489.

56. Nielsen T A. A review of mentation in REM and NREM sleep:’Covert’ REM sleep as a possible reconciliation of two opposing models. Behavioral and Brain Sciences. 2000; 23 (6): 851–86. doi: 10.1017/s0140525x0000399x.

57. Landolt H, Raimo EB, Schnierow BJ, Kelsoe JR, Rapaport MH, Gillin JC. Sleep and Sleep Electroencephalogram in Depressed Patients Treated With Phenelzine. Archives of General Psychiatry. 2001; 58 (3): 268–276. doi: 10.1001/archpsyc.58.3.268.

58. Oudiette D, Dealberto M-J, Uguccioni G, et al. Dreaming without REM sleep. Consciousness and Cognition. 2012; 21 (1): 112–117. doi: 10.1016/j.concog.2012.04.010.

59. Foulkes D. The Psychology of Sleep. 1966.

60. Fosse R, Stickgold R & Hobson J A. Thinking and hallucinating: Reciprocal changes in sleep. Psychophysiology. 2004; 41 (2): 298–305. doi: 10.1111/j.1469-8986.2003.00146.x.

61. Mamelak A N, Hobson J A. Dream bizarreness as the cognitive correlate of altered neuronal behavior in REM sleep. Journal of Cognitive Neuroscience. 1989; 1 (3): 201–22. doi: 10.1162/jocn.1989.1.3.201.

62. Hobson J A & Stickgold R. Dreaming: A neurocognitive approach. Consciousness and Cognition: An International Journal. 1994; 3 (1): 1–15. https://psycnet.apa.org/doi/10.1006/ccog.1994.1001.

63. Antrobus J. REM and NREM Sleep Reports: Comparison of Word Frequencies by Cognitive Classes. Psychophysiology. 1983; 20 (5): 562–568. 10.1111/j.14698986.1983.tb03015.x.

64. Cicogna P, Natale V, Occhionero M & Bosinelli M. Slow Wave and REM Sleep Mentation. Sleep Research Online. 2000; 3 (2): 67–72.

65. Diekelmann S & Born J. The memory function of sleep. Nature Reviews Neuroscience. 2010; 11 (2):114–126.

66. Verdone P. Temporal reference of manifest dream content. Perceptual and Motor Skills. 1965; 20(3_suppl): 1253–1268. doi: 10.2466/pms.1965.20.3c.1253.

67. Picard-Deland C, Konkoly K, Raider R, Paller K A, Nielsen T, Pigeon W R, Carr M. Thememory sources of dreams: serial awakenings across sleep stages and time of night. Sleep. 2023;46 (4). doi: 10.1093/sleep/zsac292.

68. Malinowski J E, Horton C L. Dreams reflect nocturnal cognitive processes: Early-night dreams are more continuous with waking life, and late-night dreams are more emotional and hyperassociative. Consciousness and Cognition. 2021; 88: 103071. doi: 10.1016/j.concog.2020.103071.

69. Stickgold R, Scott L, Rittenhouse C, Hobson J A. Sleep-Induced Changes in Associative Memory. Journal of Cognitive Neuroscience. 1999; 11 (2): 182–193. doi: 10.1162/089892999563319.

70. Malinowski J E. Metaphor and hyperassociativity: the imagination mechanisms behind emotion assimilation in sleep and dreaming. Frontiers in Psychology. 2015; 6: 1132. 10.3389/fpsyg.2015.01132.

71. Cicogna P, Cavallero C & Bosinelli M. Cognitive aspects of mental activity during sleep. Am J Psychol. 1991; 104 (3): 413–25.

72. Nielsen T A, Kuiken D, Alain G, Stenstrom P & Powell R. Immediate and delayed incorporations of events into dreams: further replication and implications for dream function. J. Sleep Res. 2004;13 (4): 327–36. doi: 10.1111/j.1365-2869.2004.00421.x.

73. Wamsley E J, Hirota Y, Tucker M G, Smith M R & Antrobus J S. Circadian and ultradian influences on dreaming: A dual rhythm model. Brain Research Bulletin. 2007; 71 (4): 347–354. doi:10.1016/j.brainresbull.2006.09.021.

74. Simor P, Peigneux P, & Bódizs R. Sleep and dreaming in the light of reactive and predictive homeostasis. Neuroscience and Biobehavioral Reviews. 2023; 147: 105104. doi:10.1016/j.neubiorev.2023.105104.

75. Schredl M. Laboratory references in dreams: Methodological problem and/or evidence for the continuity hypothesis of dreaming? International Journal of Dream Research. 2008; 1 (1): 3–6. doi: 10.11588/ijodr.2008.1.19.

76. Whitman R, Pierce C, Maas J & Baldrige B. The dreams of the experimental subject. Journal of Nervous and Mental Disease. 1962; 134 (5): 431–439. doi: 10.1097/00005053-196205000-00005.

77. Foulkes D & Rechtschaffen A. Presleep Determinants of Dream Content: Effects of Two Films. Perceptual and Motor Skills. 1964; 19: 983–1005. doi: 10.2466/pms.1964.19.3.983.

78. Foulkes D. Nonrapid eye movement mentation. Exp Neurol. 1967; 4: 28–38. doi: 10.1016/00144886(67)90154-9.

79. Picard-Deland C, Nielsen T & Carr M. Dreaming of the sleep lab. PLoS One. 2021; 16 (10). doi:10.1371/journal.pone.0257738.

80. Cowan E T, Schapiro A C, Dunsmoor J E & Murty V P. Memory consolidation as an adaptive process. Psychonomic Bulletin & Review. 2021; 28 (6): 1796–1810. doi: 10.3758/s13423-021-01978x.

81. Roüast N M & Schönauer M. Continuously changing memories: A framework for proactive and non linear consolidation. Trends in Neurosciences. 2023; 46 (1): 8–19. 10.1016/j.tins.2022.10.013.

82. Roenneberg T, Kuehnle T, Juda M, Kantermann T, Allebrandt K V, Gordijn M C M & Merrow M. Epidemiology of the human circadian clock. Sleep Medicine Reviews. 2007; 11 (6): 429–438. doi:10.1016/j.smrv.2007.07.005.

83. Oldfield R C. The assessment and analysis of handedness: The Edinburgh inventory. Neuropsychologia. 1971; 9 (1): 97–113. doi: 10.1016/0028-3932(71)90067-4.

84. Windt JM. Reporting dream experience: Why (not) to be skeptical about dream reports. Front Hum Neurosci. 2013; Nov 7, 7: 708. doi: 10.3389/fnhum.2013.00708.

85. Klem G H, Lüders H O, Jasper H H & Elger C. The ten-twenty electrode system of the International Federation. The International Federation of Clinical Neurophysiology. Electroencephalography and clinical neurophysiology. 1999; 52: 3–6.

86. Kumral D, Palmieri J, Gais S & Schönauer M. D. Pre-sleep experiences shape neural activity and dream content in the sleeping brain. iScience. 2025. 28, 113032 10.1016/j.isci.2025.113032.

87. Blagrove M. Distinguishing continuity/discontinuity, function, and insight when investigating dream content. International Journal of Dream Research. 2011; 4 (2): 45–47. doi: 10.11588/ijodr.2011.2.9153.

88. Schredl M & Hofmann F. Continuity between waking activities and dream activities. Consciousness and Cognition. 2003; 12 (2): 298–308. doi: 10.1016/s1053-8100(02)00072-7.

89. Kudrimoti H S, Barnes C A & McNaughton B L. Reactivation of Hippocampal Cell Assemblies: Effects of Behavioral State, Experience, and EEG Dynamics. The Journal of Neuroscience. 1999; 19 (10):4090–4101.

90. Antony J W, Cheng L Y, Brooks P P, Paller K A, Norman K A. Competitive learning modulates memory consolidation during sleep. Neurobiology of Learning and Memory. 2018; 155: 216–230. doi:10.1016/j.nlm.2018.08.007.

91. Joensen B, Harrington M O, Berens S, Cairney S A, Gaskell G & Horner A J. Targeted memory reactivation during sleep can induce forgetting of overlapping memories. Learning & Memory. 2022; 29 (10): 401–411. doi: 10.1101/lm.053594.122.

92. Picard-Deland C, Bernardi G, Genzel L, Dresler M, Schoch S F. Memory reactivations during sleep: a neural basis of dream experiences? Trends in Cognitive Sciences. 2023; 27 (6): 568–582. doi:10.1016/j.tics.2023.02.006.

